# Recovery from transgenerational RNA silencing is driven by gene-specific homeostasis

**DOI:** 10.1101/148700

**Authors:** Sindhuja Devanapally, Pravrutha Raman, Samual Allgood, Farida Ettefa, Maigane Diop, Mary Chey, Yixin Lin, Yongyi E Cho, Rui Yin, Antony M Jose

## Abstract

Changes in gene expression that last for multiple generations without changes in gene sequence have been reported in many plants and animals^1–3^. Cases of such transgenerational epigenetic inheritance (TEI) could support the ancestral origins of some diseases and drive evolutionary novelty. Here, we report that stably expressed sequences in *C. elegans* have features that provide a barrier against TEI. By using double-stranded RNA (dsRNA) targeting the same sequence in different genes, we show that genes typically recover from silencing within the germline in a few generations. A rare recombinant two-gene operon containing this target sequence that recovered poorly from induced silencing enabled us to delineate mechanisms that can perpetuate silencing. Parental exposure to dsRNA targeting one gene within this operon reveals two distinct phases of the resulting TEI: only the matching gene is silenced in early generations, but both can become silenced in later generations. However, silencing of both genes can be initiated within one generation by mating, which perturbs intergenerational RNA-based mechanisms such that silencing dominates for more than 250 generations. This stable RNA silencing can also reduce the expression of homologous sequences in different genes *in trans* within the germline, but the homologous genes recover expression after a few generations. These results suggest that stably expressed sequences are subject to feedback control that opposes TEI initiated by multiple mechanisms within the germline. We speculate that similar homeostatic mechanisms that enable recovery from epigenetic changes underlie the observed preservation of form and function in successive generations of living systems.

## Results

Changes in gene expression that persist across generations without changes in DNA sequence are easily measurable forms of transgenerational epigenetic inheritance^1–3^. Such TEI can result when a gene is silenced using RNA interference (RNAi)^4^, making it a convenient approach for inducing sequencespecific heritable change. While many studies have reported TEI occurring under diverse conditions, variation between studies precludes a consistent explanation for TEI (Extended Data Table 1). To decipher the dynamics of TEI under controlled experimental conditions, we targeted the same *gfp* sequence expressed as part of low or single-copy genes containing different regulatory sequences that all drive expression within the germline of the nematode *C. elegans.* We fed animals double-stranded RNA (dsRNA) against *gfp* and examined silencing in animals (P0) and in their untreated descendants (F1-F5) (Fig. 1a). The resulting GFP fluorescence intensity varied from bright to undetectable (“off”) among P0 animals (Extended Data Fig. 1). Out of five target genes tested with identical exposure to the initiating dsRNA, two genes showed silencing up to F2 progeny, but silencing of only one gene persisted beyond F2 (Fig. 1b, Extended Data Fig. 1). Because parental dsRNA can be deposited into progeny in *C. elegans*^5,6^, the number of generations for which ingested dsRNA can perdure is unclear. We therefore only consider changes that persist beyond the F2 generation as transgenerational silencing in this study and conclude that it is variable even when the same sequence is targeted within different genes expressed in the germline. The revival of expression in descendants despite silencing in parents suggests the presence of epigenetic recovery mechanisms that oppose change.

**Figure 1.**
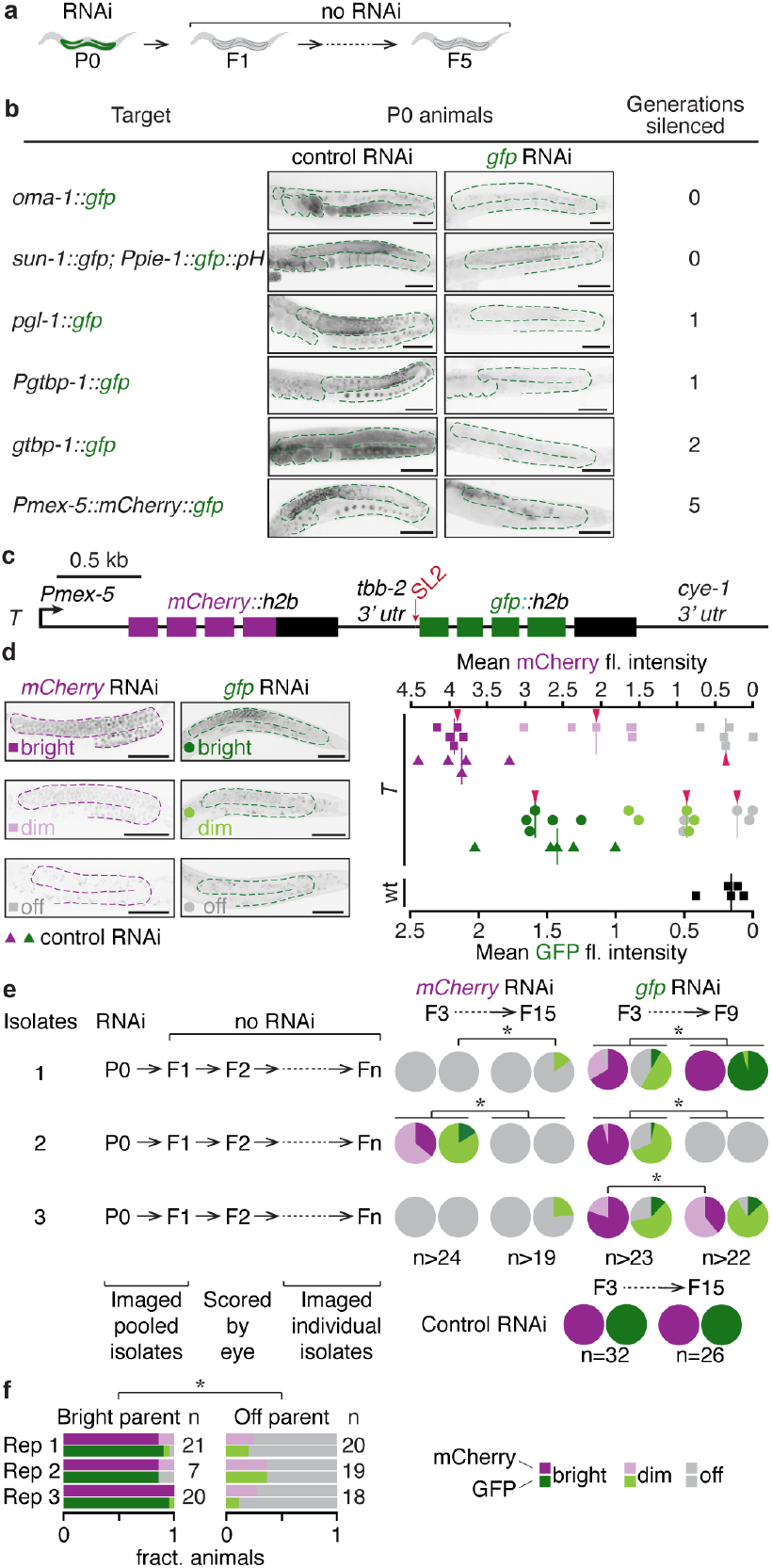
Silencing within the germline does not always initiate stable transgenerational epigenetic inheritance. **a**, Schematic of assay for transgenerational silencing. P0 animals were fed dsRNA (RNAi) for 24 hours, and the P0 animals and their untreated (no RNAi) descendants for up to five generations (F1-F5) were analysed. **b**, Five target genes containing the same *gfp* (green) sequence were exposed to the same sources of control RNAi or *gfp* RNAi. Representative images highlight the germline (green outline) of P0 animals. Numbers of descendant generations that show silencing (Generations silenced) are indicated. **c**, Schematic of the single-copy transgene *Pmex-5::mCherry::h2b::tbb-2 3’utr::gpd-2 operon::gfp::h2b::cye-1 3’ utr* called *T* in this study. d, *Left,* Representative germline images of animals expressing *T* scored as having bright (magenta or green), dim (pink or light green), or not detectable (off, grey) levels of mCherry (squares) or GFP (circles) fluorescence are shown. mCherry or GFP fluorescence within the germline was quantified in descendants of animals exposed to RNAi (control: triangles, mCherry: squares, or *gfp*: circles). *Right,* Fluorescence measured from bright, dim, off or wildtype (black squares) L4-staged hermaphrodites is plotted (n = 5). Red arrowheads correspond to animals shown on the left. e, Feeding RNAi targeting *T* was performed as in (**a**) and silencing was analysed in descendants. *Left,* All generations shown except F2s were scored by imaging. P0 and F1 were each pooled for imaging but subsequent generations each descending from one P0 ancestor were imaged as individual isolates. *Right,* Descendants of P0 ancestors exposed to *mCherry, gfp* or control RNAi were scored for expression of GFP and mCherry, and represented in a pie chart. f, Feeding RNAi targeting *T* was performed as in (**a**) by propagating twelve animals in every generation. Expression of GFP and mCherry was analysed for three replicates (Rep 1-3) in progeny of bright or off F3 animals. Asterisks indicate *P* < 0.05 using *χ*^2^ test. Scale bar (50 μm) and number of animals scored (n) are indicated. Also see Extended Data Figs. 1 and 2.

The gene^7^ that showed transgenerational silencing by feeding RNAi, hereafter referred to as *T*, can also be silenced for >25 generations by neuronal dsRNA^8^. This susceptibility to change suggests that features of *T* either recruit maintenance mechanisms or fail to recruit recovery mechanisms^9^. *T* is a single-copy transgene that encodes a bicistronic operon that expresses *mCherry* and *gfp* in the germline, presumably as one transcript before being spliced (Fig. 1c, Extended Data Fig. 2a, b). We observed transgenerational changes in GFP and mCherry expression from *T* (Fig. 1d, e) when animals were fed dsRNA against either *mCherry* or *gfp* and their descendants were propagated without bias. Upon *mCherry* RNAi, silencing of *mCherry* was observed in all generations (up to F15 tested), however, from the first generation, silencing of *gfp* was also detected, suggesting that silencing likely includes reduction of unspliced pre-mRNA from the F1 generation onwards (Fig. 1e, Extended Data Fig. 2c). In contrast, upon *gfp* RNAi, while *gfp* silencing was observed in all generations (up to F12 tested), *mCherry* silencing was robustly detectable only from the F3 generation onwards (Fig. 1e, Extended Data Fig. 2d-f). These observations suggest two distinct modes of transgenerational silencing – one that can occur without affecting pre-mRNA and another that potentially affects pre-mRNA. Similar transgenerational dynamics were observed when silenced animals were selectively propagated in every generation (Extended Data Fig. 2g) with the expression of *T* in progeny resembling parental expression (Fig. 1f). Consistent with the extreme sensitivity of *T* to TEI, feeding animals with bacteria that express a *gfp* expression vector – potentially a source of trace amounts of *gfp*-dsRNA – resulted in transgenerational silencing of *T* despite weak silencing in P0 and F1 animals (Extended Data Fig. 2h). Some studies have documented the deposition of chromatin modifications that extend to several kilobases surrounding the RNAi-targeted genomic sequence^10^ and others have suggested that chromatin modifiers are required in P0 animals^11^ for the establishment of transgenerational silencing. The transgenerational silencing of *gfp* with low *mCherry* silencing for a few generations (Fig. 1e) and in descendants without appreciable silencing in parents (Extended Data Fig. 2h) opposes the generality of these claims and suggests the existence of transgenerational silencing mechanisms that can persist with minimal need for changes in pre-mRNA or chromatin.

We found that expression of *T* in progeny depended on whether *T* was inherited paternally or maternally (Fig. 2a). This surprising difference was not observed for expression from many tested genes, including those sharing sequence identity with *T* (Extended Data Fig. 3). While progeny inheriting *T* maternally showed uniform mCherry and GFP expression, progeny inheriting *T* paternally showed loss of expression (Fig. 2a, Extended Data Fig. 4a) despite stable expression of *T* within male parents (Extended Data Fig. 2b). Hermaphrodite sperm were dispensable for this phenomenon (Extended Data Fig. 4b-d). Because this silencing can be reproducibly initiated (Fig. 2b) and is distinct from previously reported epigenetic silencing phenomena (Extended Data Table 2), we refer to it as mating-induced silencing. We systematically altered the features of *T* (Extended Data Fig. 5) and found that all tested variants were silenced (Fig. 2a, Extended Data Fig. 4e, f), suggesting that operon structure, histone sequences, *C. briggsae unc-119(+)* or the method used to insert *T* into the genome cannot explain susceptibility to mating-induced silencing. Thus, a minimal gene with *Pmex-5* driving expression of *mCherry* or *gfp* with a *cye-1* 3’ UTR (*Tcherry* or *Tgfp*) shows mating-induced silencing. Proportions of animals that showed silencing were comparable in all measured cohorts of progeny with mCherry and GFP fluorescence similarly affected within most individual F1 animals (Extended Data Fig. 4g, h), which suggests potential silencing of unspliced pre-mRNA or coordinate silencing of both *gfp* and *mCherry* mRNA after pre-mRNA splicing. Examining known RNA silencing factors^12–14^ (Extended Data Fig. 6a) revealed that mating-induced silencing required PRG-1, MUT-16, and HRDE-1 (Extended Data Fig. 6b), making it distinct from PRG-1-independent silencing by feeding RNAi (Extended Data Fig. 6c). The requirements for initiation of mating-induced silencing suggest that it relies on both small RNAs called piRNAs associated with PRG-1 and secondary small RNAs associated with HRDE-1 that are generated within perinuclear mutator foci nucleated by MUT-16^12^. The following observations support an intergenerational mechanism for the initiation of mating-induced silencing whereby maternal PRG-1-bound piRNAs trigger production of secondary small RNAs in zygotic mutator foci, which then bind HRDE-1 and are required for silencing in progeny: (i) RNA levels were reduced in silenced cross progeny (Fig. 2c, Extended Data Fig. 7a-c), (ii) removal of predicted piRNA sites^15^ in *mCherry (Tcherry-pi)* eliminated mating-induced silencing (Fig. 2d, Extended Data Fig. 4i), (iii) maternal absence of PRG-1 and zygotic absence of HRDE-1 prevented initiation (Extended Data Fig. 6d), (iv) preventing pronuclear fusion in progeny^16,17^ (Fig. 2e, f, see Methods) still resulted in silencing, indicating that maternal chromatin is not necessary in the germline for initiation.

**Figure 2.**
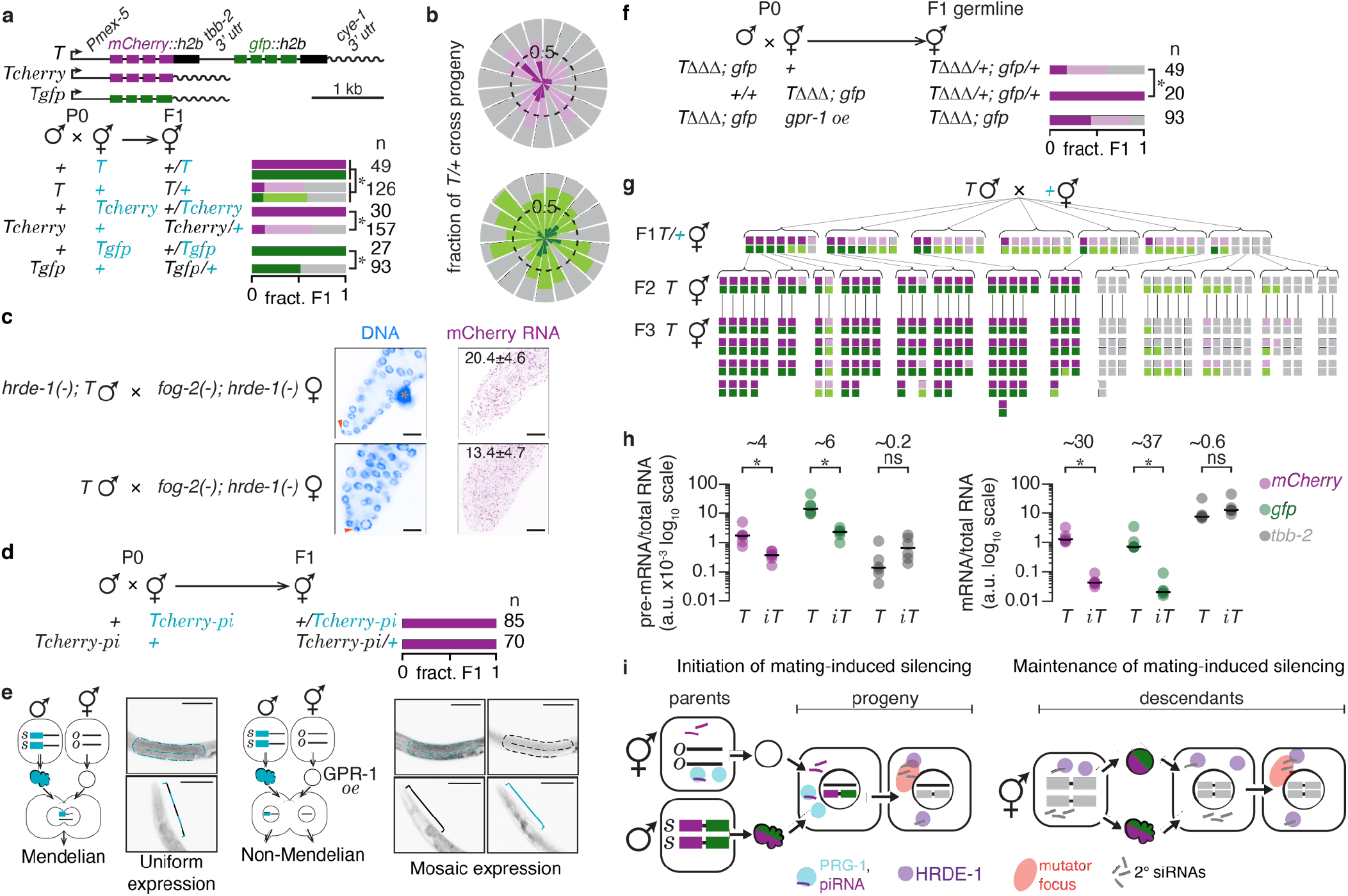
Mating can disrupt gene expression by initiating piRNA-mediated silencing. **a**, Schematics of *T* and independently generated minimal variants expressing only *mCherry* or *gfp* are depicted (*top*). Animals expressing *T, Tcherry* or *Tgfp* were mated with non-transgenic animals and resulting cross progeny were scored (*bottom*). **b**, Rose plot of independent repeats of mating-induced silencing of *T*. Each segment represents independent trials performed at different times each with up to four biological replicates and includes data from experiments depicted in other figures within the manuscript. Identically placed segments within the top and bottom plots correspond to mCherry and GFP levels obtained from the same subset of a total of 561 animals. Dashed line indicates half the fraction of animals scored. **c**, Single-molecule fluorescence *in situ* hybridization (smFISH) against *mCherry* RNA was performed in dissected gonads of animals that were impaired for (*top*) or susceptible to (*bottom*) mating-induced silencing. Images shown here are also shown in Extended Data Fig. 7 with remaining images from the same animals. Pink arrowhead, nucleus of the distal tip cell and orange asterisk, nonspecific signal (c-e). **d**, Animals expressing *Tcherry* lacking piRNA binding sites *(Tcherry-pi)* were mated with non-transgenic animals and cross progeny were scored. **e**, Scheme to test effect of *gpr-1* overexpression: *gtbp-1::gfp* (blue) males mated with wild-type hermaphrodites *(left)* or with hermaphrodites overexpressing *gpr-1* in the germline *(gpr-1 oe, right). s* and *o* label DNA inherited through sperm and oocyte respectively. Representative images show differences in segregation of *gtbp-1::gfp* in the germline (*top*) and the head (*bottom*) in cross progeny. Coloured outlines and brackets show the parental origin of germline or pharynx. Also see methods. **f**, Animals expressing *TΔΔΔ* and *gtbp-1::gfp* were mated with either non-transgenic animals or animals overexpressing *gpr-1.* Expression in the F1 germline was scored in cross progeny. **g**, Mating-induced silencing was initiated and silencing was scored in cross progeny and their descendants. Each pair of boxes represents one animal. **h**, *mCherry, gfp* and *tbb-2* pre-mRNA *(left)* or mRNA *(right)* levels were measured by qRT-PCR in animals that express *T* and in animals that showed loss of expression from *T* for >200 generations *(iT).* **i**. Model for initiation and maintenance of mating-induced silencing: PRG-1 inherited through oocyte (circle) and piRNAs are sufficient to initiate silencing of both *mCherry* and *gfp* from *T* inherited through sperm (cloud shape) into cross progeny using the secondary Argonaute, HRDE-1 and mutator proteins. Maintenance of silencing across generations requires HRDE-1 and mutator foci. Also see Methods and Extended Data Figs. 3 to 7. Asterisks indicate *P* & 0.05 and ‘ns’ indicates no significant difference using *χ*^2^ test (**a, f**) or Student’s t-test (**h**). Chromosomes with a recessive *dpy* marker (blue font), number of animals scored (n) and scale bar (50 μm) are indicated.

Once the expression state of *T* was established in cross progeny, subsequent generations tended to maintain the same expression state (Fig. 2g, Extended Data Fig. 4j). Thereafter, descendants of silenced F2 animals remained silenced for >150 generations *(iT* where *i* stands for inactive) without additional selection (Extended Data Fig. 4k-m, Extended Data Fig. 6e). Consistent with transgenerational RNA silencing, animals with *iT* showed a ~30-37 fold decrease in mRNA and ~4-6 fold decrease in pre-mRNA levels (Fig. 2h, Extended Data Fig. 7d, e). Previous studies have shown that piRNA-mediated silencing is expected to initiate stable RNA silencing leading to repressive chromatin modifications across generations^18–20^. We therefore tested if the transgenerational stability of mating-induced silencing relied on RNAi factors and found that silencing is abolished when HRDE-1 or the mutator proteins MUT-2 or MUT-16 were removed even after 250 generations of silencing (Extended Data Fig. 6e). Both maternal and zygotic HRDE-1 function together to maintain silencing (Extended Data Fig. 6f). Removal of the RNA-dependent RNA polymerases (RdRPs) EGO-1 and RRF-1, but not of RRF-1 alone, enabled a modest recovery of expression, which could imply only a modest role for small RNAs in mating-induced transgenerational silencing. However, we cannot strictly measure the need for small RNAs made by these RdRPs because maternal *ego-1* mRNA or protein could maintain silencing of *T* in progeny of *ego-1* heterozygotes (Extended Data Fig. 6e) and complete loss of EGO-1 results in sterility^21,22^. Furthermore, small RNAs made by these RdRPs do not always correlate with gene silencing^23^. Nevertheless, robust recovery of expression even after hundreds of generations of silencing suggests that silencing is actively established in every generation. Once expression is recovered in *hrde-1* mutants, restoring HRDE-1 did not re-establish silencing of *T* (Extended Data Fig. 6g), indicating that signals facilitating silencing in every generation were lost upon HRDE-1 removal. Current understanding of HRDE-1-dependent transgenerational silencing suggests that HRDE-1-bound small RNAs recognize nascent transcripts and recruit chromatin modifiers to establish repressive H3K9me3 modifications at target genes^24^. We detected no requirement for the histone methyltransferases MET-2 or SET-32^25^ or the chromodomain protein HERI-1^26^ (Extended Data Fig. 6e). Furthermore, we did not detect significant changes in H3K9 methylation (Extended Data Fig. 6h, i) in descendants from a lineage that experienced >250 generations of silencing. While TEI induced upon mating may be associated with other as yet untested molecular changes, the production of small RNAs in every generation could be sufficient for explaining the transgenerational stability of mating-induced silencing (Fig. 2i).

The stable expression of *T* observed in the absence of mating suggests that transcripts from *T* engage protective mechanisms that have been proposed to ‘license’ expression within the germline^27^. One such protective mechanism relies on phase-separated condensates within the germline called P-granules, which when disrupted can cause mis-regulation and aberrant distribution of some transcripts^28,29^. Consistent with P-granules facilitating stable expression of *T*, loss of the P-granule component PGL-1 resulted in variable expression of *T* even in the absence of mating (Extended Data Fig. 8a). Therefore, the stable expression of *T* across generations within the hermaphrodite germline reflects reliable recognition of transcripts from *T* within P-granules as part of ‘self’ in every generation^18, 30,31^.

We found that initiation of mating-induced silencing of paternally inherited *T* could be prevented by maternal expression of *T* (Fig. 3a), suggesting that maternally expressed *T* provides a separable signal that protects paternally inherited *T* from silencing. Consistently, we mapped the source of the protective signal to a ~3.2 Mb region that includes *T* (Fig. 3a). The ability to protect was also largely retained among independently generated variants of *T* (Fig. 3a, Extended Data Fig. 5, Extended Data Fig. 8b, c). Once paternally inherited *T* was protected, expression from *T* was stably maintained in descendants generated by selfing (Extended Data Fig. 8d), indicating that protection from initiation also prevents the transgenerational effects of mating-induced silencing. Nevertheless, protected cross progeny remained susceptible to initiation like unsilenced progeny that escaped initiation of mating-induced silencing (Extended Data Fig. 8e, f). Because maternally present variants of *T* with nonsense mutations or deletions could confer protection (Extended Data Fig. 8b), we examined whether the protective signal could be derived from parts of *T*. We found that *Tcherry-pi* sequences showed the strongest level of protection even when the N- or C-terminal halves of *Tcherry-pi* coding sequence were deleted (Fig. 3b), demonstrating that an identical *mCherry* coding sequence is not needed for protection and excluding the simple model of maternal piRNAs being competed away by complementary maternal *mCherry* sequences. In other words, *Tcherry-pi* can protect from mating-induced silencing despite being incapable of being silenced by the piRNAs used in mating-induced silencing. Protection was weaker with only the last exon of *Tcherry-pi* but was completely abolished when *Tcherry-pi* open reading frame was deleted (Fig. 3b). Furthermore, genes that share the same *mCherry* protein sequence or DNA sequences identical to other regions of *T* but expressed from different loci could not confer protection (Extended Data Fig. 8g, h). These findings suggest that robust protection from mating-induced silencing depends on a diffusible *mCherry* signal derived from *Tcherry(-pi).* In support of this signal being diffusible and therefore independent of direct interaction between parental chromatin for its activity, animals with impaired fusion of parental pronuclei were still protected from silencing (Extended Data Fig. 8i). Collectively, these observations suggest that protection relies on a diffusible sequence-specific signal, likely RNA. The Argonaute CSR-1 has been proposed to play a role in promoting the expression of germline genes^18,30^, although rigorous analyses are precluded by chromosome segregation defects in *csr-1* mutants that lead to embryonic lethality^32^. Furthermore, CSR-1 has been proposed to regulate spermiogenesis and oogenesis^30^, to silence sperm-specific transcripts in coordination with germ granules^33^, and to tune the levels of germline transcripts^34^. These diverse roles make effects caused by the loss of CSR-1 difficult to interpret. Nevertheless, because CSR-1-associated small RNAs have been proposed to play a role in the prevention or reversal of transgene silencing in the germline^35,36^, we examined a downstream component of the CSR-1 pathway that interacts with these small RNAs but lacks the confounding developmental defects. Unlike CSR-1, removal of the uridylyltransferase CDE-1 that uridylates CSR-1-associated small RNAs causes fewer pleiotropic effects^32,37^. CDE-1 loss did not abolish protection (Fig. 3c). Also, the protective signal could only weakly reverse silencing of *iT* (Extended Data Fig. 8j), while CSR-1-associated small RNAs were reported to robustly reverse silencing of other transgenes^36,31^. Thus, protection of *T* from mating-induced silencing relies on diffusible sequencespecific signals and could be independent of the CSR-1 pathway.

**Figure 3.**
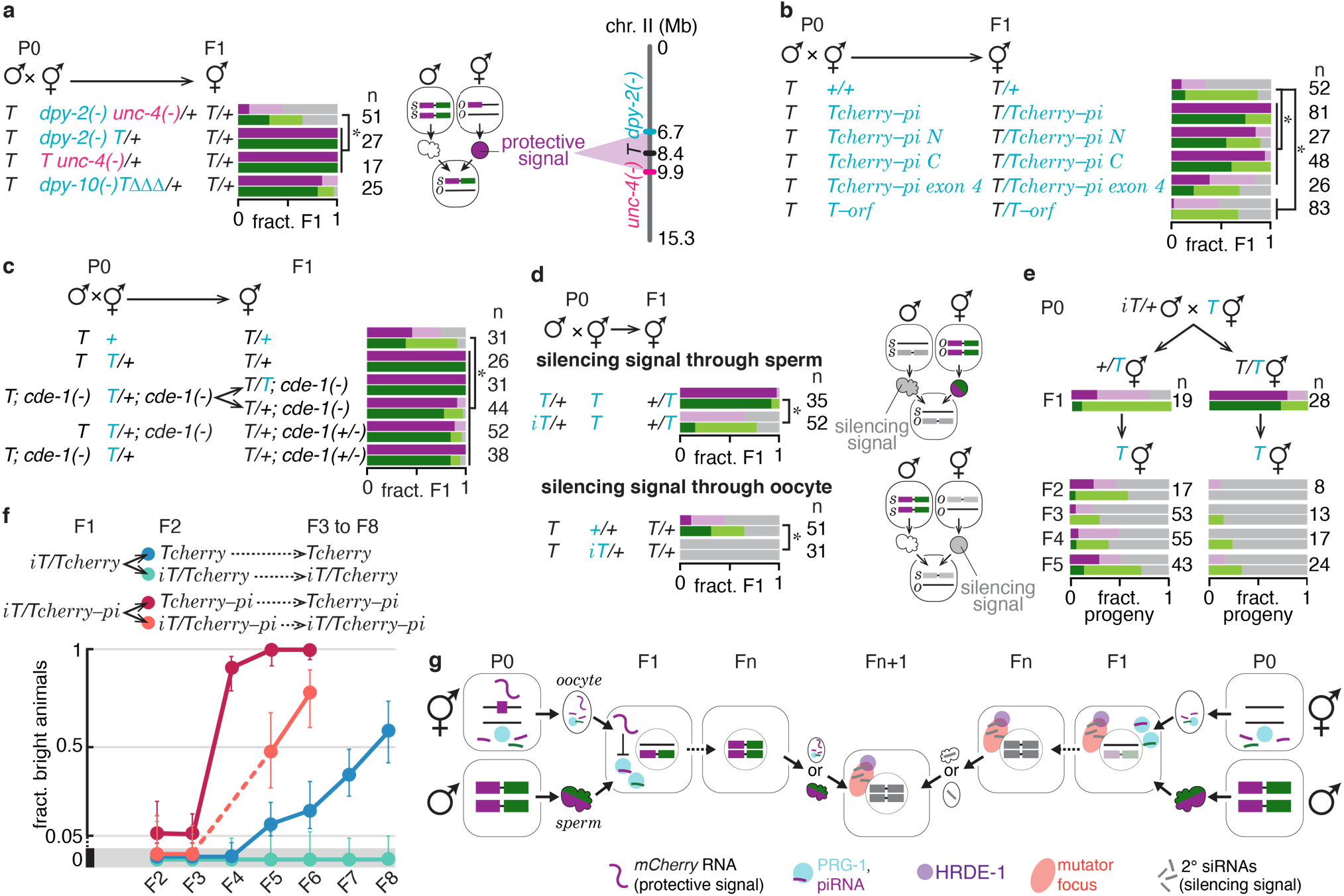
Opposing intergenerational mechanisms establish gene expression in progeny. **a**, *T* males were mated with genetically marked hermaphrodites and animals with paternally inherited *T* were scored. Schematic: maternal presence of *TΔΔΔ* protects paternally inherited *T* from mating-induced silencing, suggesting that the oocyte carries a separable protective signal derived from a region between *dpy-2* and *unc-4* that is linked to *T*. **b**, *T* males were mated with hermaphrodites expressing variants of *Tcherry-pi* and progeny with paternally inherited *T* were scored. The remaining data from this experiment are depicted in Extended Data Fig. 8c as a result of which the same control cross is displayed in both figures. **c**, Mutants of a CSR-1 pathway gene, *cde-1,* were used to test parental and zygotic requirement for protection. **d**, *T* animals were mated with non-transgenic or hemizygous *iT* animals and cross progeny that inherited only *T* were scored. Schematic: parental presence of *iT* can silence *T* inherited through the other gamete, indicating the inheritance of a separable silencing signal as schematized. **e**, Silencing of *T* by the separable silencing signal or *in trans* by *iT* was assessed across generations. **f**, *Tcherry* or *Tcherry-pi* animals were mated with *iT* stably silenced for >150 generations and fractions of animals with bright *Tcherry* or *Tcherry-pi* expression were scored in resulting cross progeny (F1) and their descendants (F3 through ≤F8). Error bars indicate 95% confidence intervals. **g**, Schematic depicts mechanisms that determine expression of *T*: maternal *mCherry* can provide a protective signal (potentially RNA) that prevents mating-induced silencing, resulting in continued expression of paternally inherited *T* in subsequent generations *(left);* parental *iT* transmits a silencing signal that uses HRDE-1-bound secondary RNAs to cause trans silencing *(right).* Also see Extended Data Figs. 5, 8 and 9. Asterisks indicate *P* < 0.05 from *χ^2^* test. Chromosomes with a recessive marker (blue or pink font), number of animals scored (n) and scale bar (50 μm) are indicated.

The stable silencing of *iT* reflects continued production of an associated silencing signal (Extended Data Fig. 8j) as revealed by two observations: (i) *iT* transmitted through one gamete could silence *T* inherited from the other gamete *in trans*, regardless of how many generations *iT* remained inactive (Extended Data Fig. 9a, b) and, (ii) presence of *iT* in one parent was sufficient to cause significant silencing of *T* inherited from the other parent (Fig. 3d). Because maintenance of *iT* requires HRDE-1 (Extended Data Fig. 6), we reasoned that this silencing *in trans* likely relies on HRDE-1-dependent small RNAs. Indeed, loss of zygotic HRDE-1 mostly eliminated *trans* silencing (Extended Data Fig. 9c). Consistent with a diffusible silencing signal, direct interaction between parental chromatin was dispensable for its activity (Extended Data Fig. 9d). This signal was not detectably inherited for more than one generation independent of *iT* and therefore depends on at least parental *iT* for stability (Extended Data Fig. 9e). Our findings implicate HRDE-1-dependent small RNAs as either the heritable silencing signal that is deposited maternally in each generation or a downstream effector that is made zygotically in each generation in response to the intergenerational silencing signal. This continuous requirement for a silencing signal is supported by recovery of expression in descendants unless *T* was continuously propagated with *iT* (Fig. 3e and Extended Data Fig. 9f). Recovery from *trans* silencing was even more robust and rapid with *Tcherry* or *Tcherry-pi* (Fig. 3f, Extended Data Fig. 9g, h), where ~60% of *Tcherry* animals and ~100% of *Tcherry-pi* animals showed recovery of complete expression within seven generations after *trans* silencing. Yet, *iT* continued to remain silenced as evidenced by absence of GFP fluorescence regardless of whether animals showed recovery of *mCherry* expression from *Tcherry* variants. These differences between *T* and *Tcherry* variants are consistent with gene-specific requirements for epigenetic recovery that oppose permanent changes in gene expression (Fig. 3g).

To evaluate the potential spread of silencing signals made by *iT*, we examined homologous sequences at other genomic positions. We observed that genes sharing coding sequence identity, but not those with only intronic or protein sequence identity, were silenced within the germline by *iT in trans* (Fig. 4a and Extended Data Fig. 10a). Such *trans* silencing of homologous loci could only be detected with a stably established *iT* but not simultaneously with initiation of mating-induced silencing of *T* (Fig. 4b). This observation suggests that the mechanism that initiates mating-induced silencing is either quantitatively distinct (e.g., increased abundance of small RNAs) or qualitatively distinct (e.g., changed timing or nature of small RNAs) from the mechanism that maintains silencing despite the shared requirement for HRDE-1 activity and mutator focus integrity. Consistent with *trans* silencing being homology-dependent, *iT*Δ established after deleting *gfp* from *T* did not silence other *gfp* genes *in trans* (Extended Data Fig. 10b). Furthermore, maternal but not paternal transmission of the silencing signal affected homologous genes, possibly reflecting differences in the nature or levels of silencing signal inherited through the two gametes (Extended Data Fig. 10c, Refs. 30,38,39). Strikingly, complete *trans* silencing of a homologous gene exhibited a switch to complete recovery within two generations (Fig. 4c), similar to recovery observed after feeding RNAi (Fig. 1b, Extended Data Fig. 1). We found that genes that recover from silencing can nevertheless require HRDE-1 for silencing (Extended Data Fig. 10d, Ref. 24). Therefore, the reason for persistent transgenerational RNA silencing versus recovery from transgenerational RNA silencing cannot be attributed solely to HRDE-1: not all HRDE-1-dependent silencing is stable. To understand the requirements for recovery, we investigated if enhancing silencing by dsRNA could inhibit recovery. Mutations in *heri-1* and *met-2* enhanced persistence of silencing (Fig. 4d, Extended Data Fig. 10e), albeit to a much lesser extent than reported in previous cases^40,41^. Similarly, removal of the endonuclease ERI-1^42^ weakly increased the persistence of silencing (Extended Data Fig. 10f, g). Nevertheless, in every case enhancing silencing still allowed recovery of resistant genes. We also detected no significant differences in abundance of RNA transcripts or subcellular localization of *T* compared to those of resistant genes (Fig. 4e, Extended Data Fig. 10h, i). Together, while most tested genes consistently recovered from transgenerational silencing and were resistant to change, *T* and its derivatives evaded epigenetic recovery and retained changes. Therefore, to understand features of a gene that enable susceptibility to mating-induced silencing we further manipulated *Tcherry. C. elegans* germline genes are under tight control of gene expression based on regulatory regions^43,44^ and on genomic position^45^ but neither altering the 3’ UTR nor changing the genomic position eliminated susceptibility of *Tcherry* to mating-induced silencing (Fig. 4f, g). Furthermore, *Tcherry* expressed from chromosome I could be protected by *Tcherry-pi* expressed from chromosome II (Fig. 4h), revealing its *trans* interaction with a nearly identical gene. Thus, the minimal gene element comprising *Tcherry* is a self-contained sequence with the ability to retain changes in expression independent of at least some genomic contexts. Underscoring the importance of gene context, the *mCherry* coding sequence from *Tcherry* is resistant to mating-induced silencing when introduced as a fusion of the endogenous *mex-5* gene (Fig. 4i). These findings suggest that *T* and its variants provide rare gene contexts that can enable coding sequences to escape recovery and retain changes in expression for many generations.

**Figure 4.**
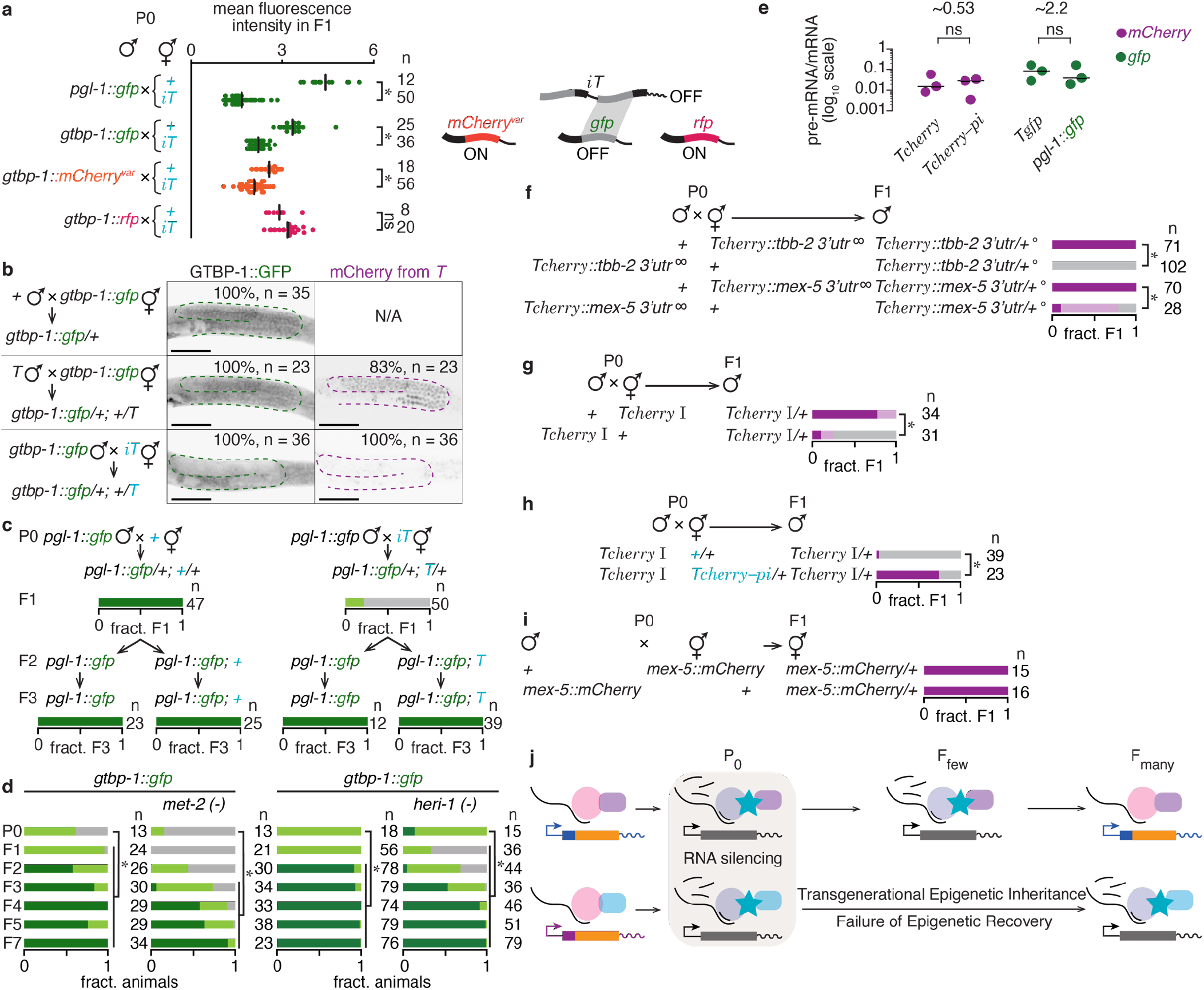
Recovery from RNA silencing is not dictated by sequence but is gene specific. **a**, Males that express homologous (*gfp*) or non-homologous (*mCherry^var^*, a synonymous *mCherry* variant or *rfp*) sequences fused to endogenous genes expressed in the germline *(pgl-1)* or ubiquitously *(gtbp-1)* were mated with non-transgenic or *iT* hermaphrodites and fluorescence of PGL-1::GFP, GTBP-1::GFP, GTBP-1::mCherry or GTBP-1::RFP was quantified in cross progeny *(left).* Schematic depicts *trans* silencing by *iT* relying on DNA sequence homology *(right).* **b**, *gtbp-1::gfp* animals were mated with non-transgenic, *T* or *iT* animals and cross progeny were imaged. Cumulative percentages of animals showing medium (representative image) or non-detectable expression level of *mCherry* from *T* are indicated. N/A, not applicable. **c**, *pgl-1::gfp* animals were mated with non-transgenic or *iT* animals and cross progeny and their descendants were scored. **d**, *gtbp-1::gfp* hermaphrodites in a wild-type, *met-2(-) (left)* or *heri-1(-) (right)* background were fed *gfp*-dsRNA for 24 hours and untreated descendants in subsequent generations (F1-F7) were scored as in Fig. 1. Feeding RNAi of other strains was performed concurrently, thus data for *gtbp-1::gfp* here is the same as in Extended Data Fig. 1c. In *heri-1(-)* animals, the statistical difference between P0 and F1/F2 is due to increased silencing, but that between P0 and F3-F7 is due to decreased silencing. Most animals fed control RNAi and descendants showed bright expression of GFP (except two out of 45 F5 descendants and one out of 37 F7 descendants of *heri-1(-)* animals that showed dim expression). **e**, pre-mRNA and mRNA levels were measured by qRT-PCR in animals expressing *mCherry* or *gfp* and depicted as a ratio. **f**, Animals expressing *Tcherry* with altered 3’ UTR were mated to non-transgenic animals and cross progeny were scored. To prevent spontaneous transgene silencing^18–20^ triggered by genome insertion, *hrde-1(-)* was introduced (∞) into P0 transgenic animals resulting in heterozygous *hrde-1(+/-)* cross progeny (°). **g-h**, *Tcherry* expressed from chromosome I was susceptible to mating-induced silencing (**g**) and protected by maternal *Tcherry-pi* (**h**). **i**, Animals with *mCherry* fused to endogenous *mex-5* gene were mated with wild-type animals and cross progeny were scored. **j**, Model depicting epigenetic recovery within the germline. Also see Extended Data Fig. 10 and Methods. Asterisks indicate *P* < 0.05 from *χ*^2^ test, ‘ns’ indicates no significant difference from *χ*^2^ test (**a**) or Student’s t-test (**e**). Chromosomes with a recessive *dpy* marker (blue font), number of animals scored (n) and scale bar (50 μm) are indicated.

We reveal that recovery mechanisms within the germline oppose transgenerational changes at the level of a gene (Fig. 4j) and maintain a transgenerational homeostasis^46^ that preserves gene expression patterns across generations. There is considerable excitement in the possibility of mechanisms that perpetuate acquired changes accelerating adaptive evolution^1,47,48^. However, indiscriminate persistence of every parental change is likely to be detrimental to organisms. Consistently, a recent measurement of changes in small RNA levels across generations in wild-type *C. elegans* suggests that such spontaneous ‘epimutations’ are maintained only for a few generations^49^. The active resistance to transgenerational epigenetic inheritance documented in this study (Fig.1, Fig. 4) suggests that organisms have evolved gene-specific mechanisms that prevent permanence of experiencedependent effects and promote recovery from epigenetic change.

## Acknowledgements

We thank Nathan Shugarts for most of the Sanger sequencing of *oxSi487,* referred to as *T* within the manuscript, presented in Extended Data Fig. 2a; members of the Jose laboratory for critical reading of the manuscript; the *Caenorhabditis elegans* Genetic Stock Center, the Seydoux laboratory (Johns Hopkins University), the Cohen-Fix laboratory (National Institutes of Health), the Fire laboratory (Stanford University), the Bringmann laboratory (Max Planck Institute) and the Hunter laboratory (Harvard University) for some worm strains. This work was supported in part by National Institutes of Health Grants R01GM111457 and R01GM124356 (to A.M.J.).

## Author contributions

All authors contributed to experimental design and analysis. S.D., P.R., S.A., F.E., M.D., Y.L, Y.E.C, M.C., and R. Y. performed experiments. S.D., P.R. and A.M.J. wrote the manuscript. All authors edited the manuscript.

## Author Information

The authors declare no competing financial interests. Correspondence and requests for materials should be addressed to A.M.J.

