## Supplementary Material for "Recovery from transgenerational RNA silencing is driven by gene-specific homeostasis"

#### Methods

##### Summary

All *C. elegans* strains were generated and maintained by using standard methods[50]. Animals with the transgene *T* (*oxSi487*) were introduced into mutant genetic backgrounds through genetic crosses using transgenic hermaphrodites and mutant males to avoid initiation of mating-induced silencing. Cross progeny from genetic crosses were identified by balancing or marking *oxSi487* with recessive mutations in *dpy-2(e8) unc-4(e120)*, *unc-4(e120)*, or *dpy-2(e8)*, *unc-8(e49) dpy-20(e1282)* and CRISPR-Cas9 generated alleles of *dpy-10* (see 'Strains used'). In some crosses, cross progeny were identified by genotyping for *oxSi487* transgene using PCR. Genome editing was performed using Cas9 protein and sgRNA[51] in most cases (Extended Data Table 3). Silencing of all transgenic strains was measured by imaging under identical non-saturating conditions using a Nikon AZ100 microscope. Quantification of images was performed using NIS Elements (Nikon) and ImageJ (NIH). Detailed procedures are provided below.

##### Strains used

| | |
|---|---|
| N2 | wild type |
| AMJ471 | *jamEx140* [*Prgef-1::gfp-dsRNA:: unc-54 3' utr & Pmyo-2::DsRed::unc-54 3' utr*] |
| AMJ501 | *oxSi487* (*Pmex-5::mCherry::h2b::tbb-2 3'utr::gpd-2 operon::gfp::h2b::cye-1 3' utr* + *unc-119(+)*) II*; unc-119(ed3)* III?; *sid-1(qt9)* V |
| AMJ506 | *prg-1(tm872)* I*; oxSi487* II; *unc-119(ed3)*? III |
| AMJ544 | *oxSi487* II*; unc-119(ed3)?* III; *nrde-3(tm1116)* X |
| AMJ545 | *oxSi487* II*; unc-119(ed3)* III?; *rde-1(ne219)* V |
| AMJ552 | *oxSi487 dpy-2(jam33)* II; *unc-119(ed3)?* III [*iT*] |
| AMJ577 | *hrde-1(tm1200)* III [4x] |
| AMJ581 | *oxSi487 dpy-2(e8)* II |
| AMJ586 | *oxSi487 dpy-2(e8)* II; *unc-119(ed3)?* III; *rde-1(ne219)* V |
| AMJ587 | *mut-2(jam9)* I |

| | |
|---|---|
| AMJ591 | jamSi25 [Punc-119deletion *jamSi19] II [TΔΔ] |
| AMJ593 | oxSi487 dpy-2(e8) II; unc-119(ed3)? III; sid-1(qt9) V |
| AMJ602 | oxSi487 dpy-2(e8) II; unc-119(ed3)? hrde-1(tm1200) III |
| AMJ626 | rrf-1(ok589) I; oxSi487 dpy-2(e8) II; unc-119(ed3)? III |
| AMJ646 | dpy-17(e164) unc-32(e189) III; rde-1(ne219) V |
| AMJ647 | dpy-17(e164) unc-32(e189) III; sid-1(qt9) V |
| AMJ667 | dpy-20(e1282) ax2053[gtbp-1::gfp] IV |
| AMJ673 | rrf-1(ok589) I; dpy-2(e8) unc-4(e120) II |
| AMJ675 | oxSi487 II; unc-119(ed3)? hrde-1(tm1200) III |
| AMJ683 | oxSi487 dpy-2(e8) II; unc-119(ed3)? III; nrde-3(tm1116) X |
| AMJ685 | K08F4.2::gfp [Pgtbp-1::gtbp-1::gfp] IV; jamEx140 |
| AMJ689 | rrf-1(ok589) I; oxSi487 II; unc-119(ed3)? III |
| AMJ690 | dpy-2(e8) unc-4(e120) II; nrde-3(tm1116) X |
| AMJ691 | dpy-2(e8) unc-4(e120) II; hrde-1(tm1200) III |
| AMJ692 | oxSi487 dpy-2(e8) II [iT] |
| AMJ693 | dpy-2(e8) unc-4(e120) II; Pmex-5::mCherry::mex-5::mex-5 3’ utr IV |
| AMJ709 | dpy-10(jam21) jamSi25 [Punc-119deletion *jamSi19] II [TΔΔ] |
| AMJ711 | prg-1(tm872) I [1x] |
| AMJ712 | dpy-2(e8) unc-4(e120) II; Pgtbp-1::gtbp-1::RFP::linker::3xflag::gtbp-1 3’utr IV |
| AMJ713 | dpy-2(e8) unc-4(e120) II; Ppgl-1::pgl-1::gfp::pgl-1 gfp 3’ utr IV |
| AMJ714 | oxSi487 II; unc-119(ed3)? hrde-1(tm1200) III |
| AMJ724 | oxSi487 II; unc-119(ed3)? III [iT] |
| AMJ725 | oxSi487 II; unc-119(ed3)? III |
| AMJ727 | dpy-2(e8) unc-4(e120) II; mCherry at cut (sens5) for gene K08F4.2 |
| AMJ753 | dpy-10(jam38) oxSi487 II; unc-119(ed3) III |
| AMJ763 | dpy-10(jam40) jamSi16 [Pmex-5::mCherry::h2b::cye-1 3’ utr *oxSi487] II [TΔ] |
| AMJ765 | dpy-10(jam41) jamSi18 [Pmex-5::mCherry::h2b::cye-1 3’ utr *oxSi487] II [TΔ] |

| | |
|---|---|
| AMJ766 | *jamSi19* [*Pmex-5::mCherry::h2b::cye-1 3' utr *oxSi487*] II [*T*Δ] |
| AMJ767 | *dpy-10(jam42) jamSi20* [*Pmex-5::mCherry::h2b::cye-1 3' utr *oxSi487*] II [*T*Δ] |
| AMJ768 | *dpy-10(jam43) jamSi21* [*Pmex-5::mCherry::h2b::cye-1 3' utr *oxSi487*] II [*T*Δ] |
| AMJ769 | *dpy-10(jam44) oxSi487* II; *unc-119(ed3)* III |
| AMJ774 | *dpy-10(jam139) jamSi23* [*Pmex-5::mCherry (6 bp indel)::h2b::tbb-2 3' utr::gpd-2 operon::gfp::h2b::cye-1 3' utr *oxSi487*] II; *unc-119(ed3)* III [*T**] |
| AMJ777 | *dpy-10(jam45)* II |
| AMJ792 | *dpy-10(jam46)* II |
| AMJ819 | *K08F4.2::gfp eri-1(mg366)* IV |
| AMJ842 | *K08F4.2::gfp eri-1(mg366)* IV; *jamEx140* |
| AMJ844 | *oxSi487 dpy-2(e8)* II [*iT*] |
| AMJ917 | *dpy-10(jam47) jamSi20* [*Pmex-5::mCherry::h2b::cye-1 3' utr *oxSi487*] II; *unc-119(ed3)* III [*iT*Δ] |
| AMJ918 | *dpy-10(jam140) jamSi32* [*Pmex-5::mCherry (3 bp indel)::h2b::cye-1 3' utr *jamSi19*] II; *unc-119(ed3)* III [*T*Δ*] |
| AMJ919 | *dpy-10(jam141) jamSi33* [*Pmex-5::mCherry (2 bp indel)::h2b::cye-1 3' utr *jamSi25*] II; *unc-119(ed3)* III [*T*ΔΔ*] |
| AMJ922 | *prg-1(tm872)* I [1x]; *dpy-2(e8) oxSi487* II; *unc-119(ed3)?* III |
| AMJ923 | *prg-1(tm872)* I [1x]; *dpy-2(e8) unc-4(e120)* II |
| AMJ926 | *dpy-10(jam39) jamSi27* [*Pmex-5::mCherry::cye-1 3' utr *jamSi25*] II [*T*ΔΔΔ] |
| AMJ928 | *jamSi27* [*Pmex-5::mCherry::cye-1 3' utr *jamSi25*] II [*T*ΔΔΔ] |
| AMJ930 | *dpy-10(jam68)* II |
| AMJ1045 | *oxSi487* II; *unc-119(ed3)? hrde-1(tm1200)* III |
| AMJ1116 | *oxSi487 dpy-2(e8)* II; *unc-119(ed3)?* III; *met-2(n4256)* III |
| AMJ1117 | *oxSi487 dpy-2(e8)* II; *unc-119(ed3)?* III; *met-2(n4256)* III |
| AMJ1118 | *oxSi487 dpy-2(e8)* II; *unc-119(ed3)?* III; *met-2(n4256)* III |
| AMJ1126 | *mut-16(pk710)* I; *oxSi487 dpy-2(e8)* II; *unc-119(ed3)?* III |

| | |
|---|---|
| AMJ1127 | *mut-16(pk710)* I; *oxSi487 dpy-2(e8)* II; *unc-119(ed3)?* III |
| AMJ1128 | *mut-16(pk710)* I; *oxSi487 dpy-2(e8)* II; *unc-119(ed3)?* III |
| AMJ1135 | *mut-2(jam9)* I; *oxSi487 dpy-2(e8)* II; *unc-119(ed3)?* III |
| AMJ1136 | *mut-2(jam9)* I; *oxSi487 dpy-2(e8)* II; *unc-119(ed3)?* III |
| AMJ1137 | *met-2(n4256)* III; *K08F4.2::gfp* IV |
| AMJ1138 | *met-2(n4256)* III; *K08F4.2::gfp* IV |
| AMJ1139 | *met-2(n4256)* III; *K08F4.2::gfp* IV |
| AMJ1142 | *oxSi487 dpy-2(e8)* II; *unc-119(ed3)?* III; *pgl-1(ct131) him-3(e1147)* IV |
| AMJ1143 | *oxSi487 dpy-2(e8)* II; *unc-119(ed3)?* III*; pgl-1(ct131) him-3(e1147)* IV |
| AMJ1157 | *oxSi487 dpy-2(jam33)* II; *unc-119(ed3)?* III; *rde-8(jam75)* IV |
| AMJ1158 | *oxSi487 dpy-10(jam82) dpy-2(jam33)* II; *unc-119(ed3)?* III; *rde-8(jam76)* IV |
| AMJ1162 | *dpy-10(jam43) oxSi487* II; *unc-119(ed3)* III |
| AMJ1170 | *jamSi37* [*Pmex-5::mCherry::cye-1 3'UTR* + *unc-119(+)*] II; *unc-119(ed3)* III |
| AMJ1174 | *dpy-10(jam106) jamSi37 [Pmex-5::mCherry::cye-1 3'UTR]* II*; unc-119(ed3)* III |
| AMJ1176 | *jamSi27* II*; K08F4.2::gfp* IV |
| AMJ1186 | *jamSi37* II*; unc-119(ed3)?* III |
| AMJ1190 | *jamSi38* [*Pmex-5::mCherry::cye-1 3'utr*] II; *unc-119(ed3)* III [$Tcherry^{Crispr}$] |
| AMJ1191 | *jamSi40* [*Pmex-5::mCherry::cye-1 3'utr*] II; *unc-119(ed3)* III [$Tcherry^{Crispr}$] |
| AMJ1192 | *jamSi41* [*Pmex-5::mCherry::cye-1 3'utr*] II; *unc-119(ed3)* III [$Tcherry^{Crispr}$] |
| AMJ1195 | *jamSi59 [Pmex-5::gfp::cye-1 3'UTR + unc-119(+)]* II*; unc-119(ed3)* III [$Tgfp$] |
| AMJ1200 | *jamSi60 [Pmex-5::gfp::cye-1 3'UTR + unc-119(+)]* II*; unc-119(ed3)* III [$Tgfp$] |
| AMJ1206 | *set-32(jam46)* I*; oxSi487 dpy-2(e8)* II*; unc-119(ed3)?* III |
| AMJ1207 | *oxSi487 dpy-2(e8) heri-1(jam47)* II*; unc-119(ed3)?* III |
| AMJ1208 | *jam148 [Pmex-5::mCherry::mex-5 3'UTR]* IV |
| AMJ1209 | *jamSi39* [*Pmex-5::mCherry (without piRNA sites)::cye-1 3' utr*] II; *unc-119(ed3)* III [$Tcherry\text{-}pi$] |

| | |
|---|---|
| AMJ1210 | *jamSi42* [*Pmex-5::mCherry (without piRNA sites)::cye-1 3' utr*] II; *unc-119(ed3)* III [*Tcherry-pi*] |
| AMJ1211 | *jamSi43* [*Pmex-5::mCherry (without piRNA sites)::cye-1 3' utr*] II; *unc-119(ed3)* III [*Tcherry-pi*] |
| AMJ1212 | *jamSi44* [*Pmex-5::mCherry (without piRNA sites)::cye-1 3' utr*] II; *unc-119(ed3)* III [*Tcherry-pi*] |
| AMJ1213 | *dpy-10(jam73) jamSi39* II; *unc-119(ed3)* III [*Tcherry-pi*] |
| AMJ1214 | *dpy-10(jam74) jamSi42* II; *unc-119(ed3)* III [*Tcherry-pi*] |
| AMJ1215 | *dpy-10(jam84) jamSi43* II; *unc-119(ed3)* III [*Tcherry-pi*] |
| AMJ1216 | *dpy-10(jam85) jamSi44* II; *unc-119(ed3)* III [*Tcherry-pi*] |
| AMJ1228 | *mut-16(pk710)* I; *oxSi487* II; *unc-119(ed3)* III |
| AMJ1236 | *jamSi37* II*; unc-119(ed3?)* III*; K08F4.2::gfp* IV |
| AMJ1238 | *dpy-10(jam106) jamSi37* II |
| AMJ1240 | *dpy-10(jam106) jamSi37* II*; ccTi1594 [mex-5p::GFP::gpr-1::smu-1 3'UTR + Cbr-unc-119(+)] unc-119(ed3?)* III |
| AMJ1245 | *jamSi61 [Pmex-5::gfp::cye-1 3' utr + unc-119(+)]* II; *unc-119(ed3)* III [*Tgfp*] |
| AMJ1248 | *dpy-10(jam142) jamSi51* [*Pmex-5::cye-1 3' utr *jamSi37*] II; *unc-119(ed3)* III [*T-orf*] |
| AMJ1249 | *dpy-10(jam143) jamSi49* [*Pmex-5::cye-1 3' utr *jamSi37*] II; *unc-119(ed3)* III [*T-orf*] |
| AMJ1259 | *hrde-1(tm1200)* III; *fog-2(q71)* V |
| AMJ1260 | *hrde-1(tm1200)* III; *fog-2(q71)* V |
| AMJ1261 | *hrde-1(tm1200)* III; *fog-2(q71)* V |
| AMJ1267 | *dpy-10(jam106) jamSi37* II*; ccTi1594 unc-119(ed3?)* III |
| AMJ1268 | *dpy-10(jam106) jamSi37* II*; ccTi1594 unc-119(ed3?)* III |
| AMJ1272 | *jamSi45* [*unc-119(+) Pmex-5::mCherry::mex-5 3' utr*] II; *hrde-1(tm1200)* III |
| AMJ1273 | *jamSi47* [*unc-119(+) Pmex-5::mCherry::mex-5 3' utr*] II; *hrde-1(tm1200)* III |
| AMJ1274 | *jamSi46* [*unc-119(+) Pmex-5::mCherry::mex-5 3' utr*] II; *hrde-1(tm1200)* III |
| AMJ1275 | *jamSi48* [*unc-119(+) Pmex-5::mCherry::mex-5 3' utr*] II; *hrde-1(tm1200)* III |

| | |
|---|---|
| AMJ1288 | *dpy-10(jam144) jamsSi52* II; *unc-119(ed3)* III [*Tcherry-pi N*] |
| AMJ1290 | *dpy-10(jam146) jamsSi54* II; *unc-119(ed3)* III [*Tcherry-pi C*] |
| AMJ1296 | *unc-119(ed3) cde-1(jam111)* III |
| AMJ1307 | *oxSi487* II*; unc-119(ed3) cde-1(jam110)* III |
| AMJ1308 | *oxSi487 dpy-10(jam138)* II*; unc-119(ed3)? cde-1(jam111)* III |
| AMJ1320 | *rrf-1(ok589) ego-1(jam93)* I |
| AMJ1321 | *rrf-1(ok589) ego-1(jam93)* I |
| AMJ1336 | *dpy-10(jam147) jamSi57* [*Pmex-5::mCherry(exon 4)::cye-1 3' utr *jamSi39*] II; *unc-119(ed3)* III [*Tcherry-pi exon 4*] |
| AMJ1337 | *dpy-10(jam149) jamSi58* [*Pmex-5::mCherry(exon 4)::cye-1 3' utr *jamSi39*] II; *unc-119(ed3)* III [*Tcherry-pi exon 4*] |
| AMJ1338 | *jamSi56* II; *unc-119(ed3)* III [*Tcherry* I] |
| AMJ1339 | *jamSi63* [*unc-119(+) Pmex-5::mCherry::tbb-2 3' utr*] II; *hrde-1(tm1200)* III |
| AMJ1340 | *jamSi64* [*unc-119(+) Pmex-5::mCherry::tbb-2 3' utr*] II; *hrde-1(tm1200)* III |
| AMJ1341 | *jamSi65* [*unc-119(+) Pmex-5::mCherry::tbb-2 3' utr*] II; *hrde-1(tm1200)* III |
| DR439 | *unc-8(e49) dpy-20(e1282)* IV |
| EG4322 | *ttTi5605* II; *unc-119(ed9)* III |
| EG6787 | *oxSi487* II*; unc-119(ed3)* III |
| EG6771 | *oxSi466* [*Pdpy-30::gfp::h2b::tbb-2 cb-unc-119(+)*] II; *unc-119(ed3)* III [gift from Christian Frøkjær-Jensen] |
| EG6779 | *oxSi474* [*Pdpy-30::gfp::h2b::tbb-2 cb-unc-119(+)*] I; *unc-119(ed3)* III [gift from Christian Frøkjær-Jensen] |
| EG6808 | *unc-119(ed3)* III; *oxTi132* [*Pdpy-30::gfp::h2b::tbb-2 cb-unc-119(+)*] V (*him-5* in background?) [gift from Christian Frøkjær-Jensen] |
| EG6810 | *unc-119(ed3)* III; *oxTi134* [*Pdpy-30::gfp::h2b::tbb-2 cb-unc-119(+)*] I (*him-5* in background?) [gift from Christian Frøkjær-Jensen] |

| | |
|---|---|
| EG6814 | *unc-119(ed3)* III; *oxTi138* [*Pdpy-30::gfp::h2b::tbb-2 cb-unc-119(+)*] I (*him-5* in background?) [gift from Christian Frøkjær-Jensen] |
| EG6838 | *unc-119(ed3) oxTi162* [*Pdpy-30::gfp::h2b::tbb-2 cb-unc-119(+)*] III (*him-5* in background?) [gift from Christian Frøkjær-Jensen] |
| GE1708 | *dpy-2(e8) unc-4(e120)* II |
| GR1373 | *eri-1(mg366)* IV |
| HC196 | *sid-1(qt9)* V |
| HC780 | *rrf-1(ok589)* I |
| HT1593 | *unc-119(ed3)* III |
| JH3197 | *ax2053* (*gtbp-1::gfp*) IV [gift from Geraldine Seydoux] |
| JH3270 | *Ppgl-1::pgl-1::gfp::pgl-1 gfp 3' utr* IV [gift from Geraldine Seydoux] |
| JH3296 | *Pmex-5::mCherry::mex-5::mex-5 3' utr* IV [gift from Geraldine Seydoux] |
| JH3323 | *Pgtbp-1::gtbp-1::mCherry::gtbp-1 3' utr* IV [gift from Geraldine Seydoux] |
| JH3337 | *Pgtbp-1::gtbp-1::RFP::linker::3xflag::gtbp-1 3'utr* II [gift from Geraldine Seydoux] |
| MT13293 | *met-2(n4256)* III |
| NL1810 | *mut-16(pk710)* I |
| OCF62 | *jfSi1* [*Psun-1::gfp cb-unc-119(+)*] II*; ltIs38* [*(pAA1) pie-1::GFP::PH(PLC1delta1) + unc-119(+)*] [gift from Orna Cohen-Fix] |
| OCF69 | *ocfSi1* [*Pmex-5::Dendra2::his-58::tbb-2 3' utr + unc-119(+)*] I*; unc-119(ed3)* III [gift from Orna Cohen-Fix] |
| PD1594 | *ccTi1594 unc-119(ed3)* III (*gpr-1 oe*) |
| SP471 | *dpy-17(e164) unc-32(e189)* III |
| SS2 | *pgl-1(ct131) him-3(e1147)* IV |
| TX189 | *unc-199(ed3)* III*; teIs1* [*(pRL475) oma-1p::oma-1::GFP + (pDPMM016) unc-119(+)*] IV |
| WM27 | *rde-1(ne219)* V |
| WM156 | *nrde-3(tm1116)* X |
| WM161 | *prg-1(tm872)* I |

All strains with fluorescent reporters showed invariable expression of fluorescence, except OCF69 which showed suppression of expression in one of the 34 tested animals.

**Primers, smFISH probes and CRISPR sequences used**

P1 ATAAGGAGTTCCACGCCCAG

P2 CTAGTGAGTCGTATTATAAGTG

P3 TGAAGACGACGAGCCACTTG

P4 ATCGTGGACGTGGTGGTTAC

P5 CTCATCAAGCCGCAGAAAGAG

P6 GGTTCTTGACAGTCCGAACG

P7 ACGGTGAGGAAGGAAAGGAG

P8 ACAAGAATTGGGACAACTCCAG

P9 AGTAACAGTTTCAAATGGCCG

P10 TCTTCACTGTACAATGTGACG

P11 CACTATTCACAAGCATTGGC

P12 CGGACAGAGGAAGAAATGC

P13 TGCCATCGCAGATAGTCC

P14 TGGAAGCAGCTAGGAACAG

P15 CCGTGACAACAGACATTCAATC

P16 ACGATCAGCGATGAAGGAG

P17 GGAGATCCATGATTAGTTGTGC

P18 GCAGGCATTGAGCTTGAC

P19 TCATCTCGGTACCTGTCGTTG

P20 AGAGGCGGATACGGAAGAAG

P21 CATAACCGTCGCTTGGCAC

P22 TCGAGTCGTGGTACAGATCG

P23 CATGCTCGTCGTAATGCTCG

P24 CGATCGTGCCAGAACAATCC

P25 ATGAAAGCCGAGCAACAACG

P26 AGAATGATGAGTCGCCACAGG

P27 CATGCACAACAAAGCCGACTAC

P28 TGAGAATACGGTCGCAGTTAGG

P29 ACGGATGCCTAGTTGCATTG

P30 CCTTCCCAGAGGGATTCAAGTG

P31 TCTGTTCCTATTCTGTCTGCAC

P32 CGCGGTTCGCAATAGGTTTC

P33 TCACCTAGTCTGTGCCATTTC

P34 TGCGGGTTTCTGTTAGCTTC

P35 GCACAGACTAGGTGAAAGAGAG

P36 ACCTCCCACAACGAGGATTAC

P37 TGGGCGTGGAACTCCTTATC

P38 GGCGAAGAGCAAAGCAGAG

P39 GGGCCGTTATCCTTTCAAATGC

P40 CATGGGCCACGGATTGTAAC

P41 ACGCATCTGTGCGGTATTTC

P42 ATTTAGGTGACACTATAGGATCAGGTAGTGGCCCACCAGTTTTAGAGCTAGA AATAGCAAG

P43 AAAAGCACCGACTCGGT

P44 ATGGTCTCCAAGGGAGAGGAG

P45 GAATCCTATTGCGGGTTATTTTAGCCACTACCTGATCCCTTG

P46 ATTTAGGTGACACTATAGGTGTAATCCTCGTTGTGGGGTTTTAGAGCTAGAAATAGCAAG

P47 CAAGGGATCAGGTAGTGGCTAAAATAACCCGCAATAGGATTC

P48 TAAGGAGTTCCACGCCCAG

P49 TTTCGCTGTCCTGTCACACTC

P50 CGATGATAAAAGAATCCTATTGCGGGTTATTTTTTGAGCCTGCTTTTTTGTACAAACTTG

P51 CAAGTTTGTACAAAAAAGCAGGCTCAAAAAATAACCCGCAATAGGATTCTTTTATCATCG

P52 AGCTAACAGAAACCCGCATAC

P53 CCTGTCACACTCGCTAAAAACAC

P54 ACAGAAACCCGCATACTCG

P55 ATTTAGGTGACACTATAGATTCCTTGTTCGGTGCTTGGGTTTTAGAGCTAGAAATAGCAAG

P56 ATTCCATGATGGTAGCAAACTCACTTCGTGGGTTTTCACAACGGCAAAATATCAGTTTTT

P57 ATTTAGGTGACACTATAGCTACCATAGGCACCACGAGGTTTTAGAGCTAGAAATAGCAAG

P58 CACTTGAACTTCAATACGGCAAGATGAGAATGACTGGAAACCGTACCGCATGCGGTGCCTA
TGGTAGCGGAGCTTCACATGGCTTCAGACCAACAGCCTA

P59 ATTTAGGTGACACTATAGACAAATGCCCGGGGGATCGGGTTTTAGAGCTAGAAATAGCAAG

P60 TGAGGTCAAGACCACCTACAAG

P61 GAATCCTATTGCGGGTTATTTTACTTGCTGGAAGTGTACTTGG

P62 CCAAGTACACTTCCAGCAAGTAAAATAACCCGCAATAGGATTC

P63 GACCACCTACAAGGCTAAGAAG

P64 ATTTAGGTGACACTATAGGGGAGAGGGAAGACCATACGGTTTTAGAGCTAGAAATAGCAAG

P65 GCAAAAATTCCCCGACTTTCCC

P66 GAAAAGTTCTTCTCCTTTACTCATTTTTGAGCCTGCTTTTTTGTAC

P67 GTACAAAAAAGCAGGCTCAAAAATGAGTAAAGGAGAAGAACTTTTC

P68 CCCATGGAACAGGTAGTTTTCC

P69 CGACTTTCCCCAAAATCCTGC

P70 ACAGGTAGTTTTCCAGTAGTGC

P71 AGAGGGATTCAAGTGGGAGAG

P72 TGGGTCTTACCGCGTATACC

P73 TGATCCCTTGTAAAGCTCATCC

P74 GTGTGTGCTGCTCGGTTAAG

P75 AATTCCACAGTTGCTCCGAC

P76 TCATCTCGCCCGATTCATTG

P77 CCGTTTCTTCCTGGTAATCC

| | |
|---|---|
| P78 | GGGTGAAGGTGATGCAACATAC |
| P79 | GGGACAACCTGTGTGCATG |
| P80 | AAGGTCCACATGGAGGGATC |
| P81 | AAAGTAATTCTACAGTATTCCTGAGATG |
| P82 | CGTCTCTTGATATTCCTTGC |
| P83 | CCAAGCGAATGGAAGCTGAAAATT |
| P84 | CAAGCGAATGGAAGTGGTCCT |
| P85 | GTAGTGACAAGTGTTGGCCATGG |
| P86 | TCACATACACATCTTCTGCACC |
| P87 | TTGGTAGAAGCTGCATCACTTT |
| P88 | CCAGACGGAACCTTCAAG |
| P89 | TCCGTCTGAAAAAATTTAATTAATT |
| P90 | GAGATTCAAGGTCCACATGGAGG |
| P91 | ATGGAAGTGGTCCTCCCTTGG |
| P92 | TCTTCGGCGCTAATCTTTTC |
| P93 | CACGAGTTCGAGATCGAG |
| P94 | GTCATCTCCGACGAGCAC |
| P95 | TTCCGTTGTTGGCTTCGTTG |
| P96 | GAGATTCAAGGTCCACATGGAGG |
| P97 | ATGGAAGTGGTCCTCCCTTGG |
| P98 | GGTGATGTTAATGGGCAC |
| P99 | TGTTGGCCATGGAACAGG |
| P100 | ATTTAGGTGACACTATAGGATTACTCATAATGACATGGTTTTAGAGCTAGAAATAGCAAG |
| P101 | GGACCACGTGGAGTTCCAGGACATCCAGGTTTTCCAGGTGACCCAGGAGAGTATGGAATT |
| P102 | ATTTAGGTGACACTATAGCGTTGGTGATGGTGATGAGGTTTTAGAGCTAGAAATAGCAAG |
| P103 | ATCTGATTATTATATTTCAGATTACTCATAATTAATGTATTCAATTTGTTAATATATTTC |
| P104 | ATTTAGGTGACACTATAGTGCTTCGATAGATCTCGAGGTTTTAGAGCTAGAAATAGCAAG |

P105 ATTTAGGTGACACTATAGTTCAGCTTACAATGGACTAGTTTTAGAGCTAGAAATAGCAAG

P106 TTAATTCTTAACAAAAAACTGTTTCCGCTCCTACGGATACAACTACATGAAAAATCATCT

P107 ATTTAGGTGACACTATAGAGTAGTTACTGATGAGCTGGTTTTAGAGCTAGAAATAGCAAG

P108 ATTTAGGTGACACTATAGTCGAGCTGTAGGCTCTTGGGTTTTAGAGCTAGAAATAGCAAG

P109 GAGAGATTCAAAAGAACAAAAAAGCCGCAGAGAGCCTACAGCTCGATCTGTAGAGTGTTT

P110 GCUACCAUAGGCACCACGAGGUUUUAGAGCUAUGCU

P111 AGCAUAGCAAGUUAAAAUAAGGCUAGUCCGUUAUCAACUUGAAAAAGUGGCACCGAGU
CGGUGCUUU

P112 TGATGATAGCCATGTTATCC

P113 GTGGACCTTGAATCTCATGA

P114 CTCTCCCTCGATCTCGAACTCGTGTC

P115 CTTGGTGACCTTAAGCTTAG

P116 GATATCCCAAGCGAATGGAA

P117 CGTACATGAACTGTGGGGAA

P118 TGCTTGACGTAAGCCTTGGA

P119 GGTAATCTGGGATATCAGCT

P120 GAATCCCTCTGGGAAGGAAA

P121 ATCCTCGAAGTTCATGACTC

P122 GAATCCTGGGTGACGGTGAC

P123 ATGAACTCTCCATCCTGAAG

P124 TCCTCTAAGCTTGACCTTGT

P125 GTCCATCGGATGGGAAGTTG

P126 ATGGTCTTCTTCTGCATGAC

P127 TACATTCTCTCGGAGGAAGC

P128 CTTGATCTCTCCCTTAAGAG

P129 TCCATCCTTAAGCTTAAGTC

P130 TTGACCTCAGCATCGTAGTG

| | |
|---|---|
| P131 | CTTCTTAGCCTTGTAGGTGG |
| P132 | TAAGCTCCTGGAAGCTGGAC |
| P133 | ATCAAGCTTGATGTTGACGT |
| P134 | TGTAATCCTCGTTGTGGGAG |
| P135 | CTCTCGTACTGCTCGACGAT |
| P136 | TTGTAAAGCTCATCCATTCC |
| P137 | AAGTTCTTCTCCTTTACTCA |
| P138 | GAATTGGGACAACTCCAGTG |
| P139 | CCCATTAACATCACCATCTA |
| P140 | CCTCTCCACTGACAGAAAAT |
| P141 | GTAAGTTTTCCGTATGTTGC |
| P142 | TGGAACAGGTAGTTTTCCAG |
| P143 | GGTATCTCGAGAAGCATTGA |
| P144 | TCATGCCGTTTCATATGATC |
| P145 | GGGCATGGCACTCTTGAAAA |
| P146 | TTCTTTCCTGTACATAACCT |
| P147 | GTTCCCGTCATCTTTGAAAA |
| P148 | CCTTCAAACTTGACTTCAGC |
| P149 | ACCTTTTAACTCGATTCTAT |
| P150 | GTGTCCAAGAATGTTTCCAT |
| P151 | GTGAGTTATAGTTGTATTCC |
| P152 | GTCTGCCATGATGTATACAT |
| P153 | CTTTGATTCCATTCTTTTG |
| P154 | CCATCTTCAATGTTGTGTCT |
| P155 | ATGGTCTGCTAGTTGAACGC |
| P156 | CGCC AATTGGAGTA TTTTGT |
| P157 | GTCTGGTAAAAGGACAGGGC |

P158 AAGGGCAGATTGTGTGGACA

P159 TCTTTTCGTTGGGATCTTTC

P160 TCAAGAAGGACCATGTGGTC

P161 AATCCCAGCAGCTG TTACAA

P162 TATAGTTCATCCATGCCATG

P163 ATTTAGGTGACACTATAGTCAACTTCTAATTTTAATTCGTTTTAGAGCTAGAAATAGCAAG

P164 ATTTAGGTGACACTATAGGTGATGAACTTCGAGGATGGGTTTTAGAGCTAGAAATAGCAAG

P165 ATTTAGGTGACACTATAGCTTTACAAGGGATCAGGTAGGTTTTAGAGCTAGAAATAGCAAG

P166 ATTTAGGTGACACTATAGAAAAATGGTCTCCAAGGGAGGTTTTAGAGCTAGAAATAGCAAG

P167 ATTTAGGTGACACTATAGCCTTCCCAGAGGGATTCAAGGTTTTAGAGCTAGAAATAGCAAG

P168 TCTCCTTCCCAGAGGGATTCAAGTGGGAGAGAGTGTAAAATAACCCGCAATAGGATTCTTT TATCATCGA

P169 CAGAGACAAGTTTGTACAAAAAAGCAGGCTCAAAAATGAACTTCGAGGAT GGAGGAGTCGTCACCGTCAC

P170 ATTTAGGTGACACTATAGAATGGTCTCCAAGGGAGAGGGTTTTAGAGCTAGAAATAGCAA G

P171 ATTTAGGTGACACTATAGCTTTACAAGGGATCAGGTAGGTTTTAGAGCTAGAAATAGCAAG

P172 CAGAGACAAGTTTGTACAAAAAAGCAGGCTCAAAAAATAACCCGCAATAGGATTCTTTTATC ATCGAAAT

P173 ATTTAGGTGACACTATAGAAAAATGGTCTCCAAGGGAGGTTTTAGAGCTAGAAATAGCAA G

P174 ATTTAGGTGACACTATAGTAATCTGATTTAAATTTTCAGTTTTAGAGCTAGAAATAGCAAG

P175 AGACAAGTTTGTACAAAAAAGCAGGCTCAAAAATGGGACACTACGATGCTGAGGTCAAGAC CACCTACAA

**Nomenclature of transgenes**

The letter *T* is used to specify the transgene *oxSi487* in all genetic crosses. The active or expressing allele of *oxSi487* is named as *T* and the inactive or the silenced allele of *oxSi487* is named as *iT* in parents. Genotypes that additionally include a recessive marker (*dpy* or *dpy unc*) are in blue or pink font.

See Extended Data Fig. 5 for all variants of *T* and 'Genetic Crosses' for details on recessive mutations used.

**Feeding RNAi and scoring associated defects**

RNAi experiments were performed at 20°C on nematode growth media plates supplemented with 1 mM IPTG (Omega Bio-Tek) and 25g/ml Carbenicillin (MP Biochemicals) (RNAi plates). In all cases genotype- and age-matched animals were fed control RNAi (L4440) and scored alongside as a control.

*Single generation (P0 Feeding RNAi)*

This assay was performed as described previously[5] and was used in all figures with feeding RNAi except Extended Data Fig. 10. Briefly, L4 animals were fed dsRNA against target genes for 24 hours. Some P0 animals were scored for expression while remaining were washed four times in M9 buffer and then allowed to crawl on unseeded plates for an hour to get rid of residual RNAi food. Animals were then singly placed on OP50 and 6 to 12 L4 animals were blindly passaged every 3 to 4 days to prevent starvation and to keep track of the generations post feeding. L4 animals were scored in each generation by imaging and L4 siblings were passaged to obtain progeny for the next generation. In feeds performed in Extended data Fig. 2d, e, F2 animals were scored by eye as noted in the schematic in Fig. 1e.

*Multiple generations (P0-F2 Feeding RNAi)*

Multiple generations of animals (P0-F2) were subjected to feeding RNAi. F1 and F2 animals were scored at L4 stage to assess the potency of the RNAi food and L4 stage siblings were transferred to a new plate with RNAi food to prevent starvation. Similar to the P0 Feeding RNAi protocol, adults (24 hours post L4) were washed four times with M9 buffer to remove residual dsRNA and transferred to a plate with OP50. Untreated progeny were then scored for inherited silencing effects. This assay was used in Extended Data Fig. 10g.

**Expression of dsRNA**

To study inherited silencing, we expressed dsRNA from an extrachromosomal array that is mitotically unstable. Animals that express the array will have both progeny that inherit the array and those that do not. We used an array expressing dsRNA in neurons and DsRed in the pharynx from *jamEx140 [Prgef-1::gfp-dsRNA:: unc-54 3'UTR & Pmyo-2::DsRed::unc-54 3'UTR]*[8]. Progeny that lack the array were evaluated to measure inherited silencing since parents were exposed to dsRNA from the array but progeny were not. This assay was used in Extended Data Fig. 10f.

**Quantification of silencing and measurement of fluorescence intensity**

To classify fluorescence intensity, in most cases, animals of the L4 stage or 24 hours after the L4 stage were mounted on a slide after paralyzing the worm using 3 mM levamisole (Sigma-Aldrich, Cat# 196142), imaged under non-saturating conditions (Nikon AZ100 microscope and Photometrics Cool SNAP $HQ^2$ camera), and binned into three groups – bright, dim and off. A C-HGFI Intensilight Hg Illuminator was used to excite GFP or Dendra2 (filter cube: 450 to 490 nm excitation, 495 dichroic, and 500 to 550 nm emission) or mCherry or RFP (filter cube: 530 to 560 nm excitation, 570 dichroic, and 590 to 650 nm emission). Sections of the gonad that are not obscured by autofluorescence from the intestine were examined to classify GFP and mCherry fluorescence from *oxSi487*. Autofluorescence was appreciable when imaging GFP but not when imaging mCherry. In some cases, fluorescence intensity within the germline was scored by eye at L4 stage (Extended Data Fig. 10 f, g) or at 24 hours after the L4 stage (Extended Data Figs. 2d, e (F2 animals only) and Extended Data Figs. 4j, k) at fixed magnification and zoom using the Olympus MVX10 fluorescent microscope without imaging.

To quantitatively measure fluorescence of mCherry from *T* (Fig. 1, Extended Data Fig. 4a) and fluorescence from other transgenes (Fig. 4a, Extended Data Fig. 1), regions of interest (ROI) were marked using either NIS elements or ImageJ (NIH) and the intensity was measured. Background was subtracted from the measured intensity for each image. For Fig. 1, Extended Data Fig. 1, Extended Data Fig. 4 and Fig. 4, fluorescence intensity was measured as x-b, where x = mean intensity of ROI and b = mean intensity of background. The obtained intensity values were converted to a $\log_2$ scale and plotted. In experiments with feeding RNAi, target gene (*gfp* or *mCherry*) and control RNAi fed animals for each strain were imaged at the same exposure. Control and experimental animals were all imaged at non-saturating conditions either at a fixed exposure (GFP-filter cube: 450 to 490 nm excitation, 495 dichroic, and 500 to 550 nm emission or mCherry-filter cube: 530 to 560 nm excitation, 570 dichroic, and 590 to 650 nm emission) or by setting exposure to their respective controls. Previous reports have suggested that the pharynx, neurons, and vulval muscles can be resistant to silencing[4,52] by dsRNA and hence were not included in our scoring.

All images being compared were adjusted identically using Adobe Photoshop for display.

*Quantification of expression from Tgfp*

Insertion of *Tgfp* into the genome resulted in variable GFP expression in all animals. However, in the case of mating-induced silencing, silenced animals displayed no detectable silencing of GFP as measured by quantification. To quantitatively measure fluorescence of GFP from *Tgfp* (Extended Data Fig. 4e), ROI of the germline that excluded the intestine was marked using Fiji (NIH) and the intensity was measured. An area outside the worm within the same image was measured for background intensity. The mean fluorescence intensity from *Tgfp* expression was calculated by subtracting the background intensity from measured GFP intensity.

**Stages of worms that were imaged**

Fluorescence intensity of mCherry or GFP was scored in L4-staged animals in all feeding RNAi experiments except in P0 RNAi fed animals, animals expressing *oma-1::gfp* or *Ppie-1::gfp::pH* (Fig 1, Extended Data Figs. 1, 2). Fluorescence intensity of mCherry or GFP was scored in L4-staged animals represented in Fig. 1d, e, Extended Data Fig. 1a-e, Extended Data Fig. 2b-h, Fig. 2a, b, d, g, Extended Data Fig. 3, Extended Data Fig. 4a-i, k-m, Extended Data Fig. 6b, d-g, Extended Data Fig. 7a, Fig. 3a, b, d, f, Extended Data Fig. 8a-h, i, Extended Data Fig. 9a-c, e, g, h, Fig. 4a, b, c, i., Extended Data Fig. 10a-g. Fluorescence intensity of mCherry or GFP was scored in adults at 24 hours post L4 stage in P0 animals represented in Fig. 1b, e, f, Extended Data Fig. 1a-e, Extended Data Fig. 2c-h, Fig. 4d, Extended Data Fig. 4d, e and all animals represented in Fig. 2b, Extended Data Fig. 3, Extended Data Fig. 4j, Extended Data Fig. 6f, Fig. 3d, e, Extended Data Fig. 8d, i, Extended Data Fig. 9e, f, Fig. 4 f-h.

**Genetic Crosses**

Three L4 hermaphrodites and 7-13 males were placed on the same plate and allowed to mate in each cross plate. Cross progeny were analyzed three to five days after the cross plate was set up. At least two independent matings were set up for each cross. For crosses in Extended Data Fig. 4k, j, the required genotypes were determined by PCR (primers P1, P2, and P3) after scoring all animals and only the data from animals with the correct genotypes were plotted. In Fig. 2b, Extended Data Fig. 3b, Extended Data Fig. 4a, b, c, g, h, Extended Data Fig. 6b, e, f, g, i, Fig. 3a, d, e, Extended Data Fig. 8a, d-g, i, Extended Data Fig. 9a, b, c, e, f, Fig. 4a, b, c, and Extended Data Fig. 10a, b, c, *dpy-2(e8)* (~3 cM from *oxSi487*) was used as a linked marker or balancer to determine the genotype of *T*. In Fig. 2a, b, d, g, Extended Data Fig. 4e, f, i, l, m Fig. 3a, b, f, Extended Data Fig. 8b-d, and Extended Data Fig. 9g, h, Fig. 4h, *dpy-10(-)* (~7 cM from *oxSi487*) was used as a linked marker or balancer to determine the genotype of *T.* In Extended Data Fig. 3b, Fig. 4b, Extended Data Fig. 8h, Extended Data Fig. 10c, *unc-8(e49) dpy-20(e1282)* was used as a linked marker or balancer to determine the genotype of *ax2053*. In Fig. 3a, *unc-4(e120)* (~1.5 cM from *oxSi487*) was used as a linked marker or balancer to determine the genotype of *T.* In Extended Data Figs. 6b *right* (control for *rde-1(-)*), *dpy-17(e164) unc-32(e189)* were used as markers to facilitate identification of cross progeny. Some crosses additionally required identification of cross progeny by genotyping of single worms, including those from Fig. 3a, d, e, Extended Data Fig. 6e (for *ego-1(-) rrf-1(-)*), Extended Data Fig. 6g, Extended Data Fig. 8c, h and Extended Data Fig. 10b, c. Animals from crosses with *prg-1(+/-)* males in Extended Data Figs. 6b *right* and 6f or with *T; prg-1(+/-)* males in Extended Data Figs. 6b *left* were also genotyped to identify *T; prg-1(-/-)* or *prg-1(-/-)* cross progeny, respectively. In crosses from Extended Data Figs. 8f and 10b, cross progeny of the required genotype were identified by the absence or presence of pharyngeal mCherry or GFP[8], respectively.

*Genetic crosses with* mut-16 *mutants to test for initiation of mating-induced silencing*

In Extended Data Fig. 6b, L4 male cross progeny were scored for only mCherry fluorescence because GFP fluorescence was difficult to assess in the single gonad arm of the L4 male germline due to gut autofluorescence.

*Genetic crosses to determine if recovery of expression upon removal of hrde-1 is lost upon re-introduction of hrde-1*

In Extended Data Fig. 6g, *hrde-1(-)* mutant males were mated with $iT$ hermaphrodites that remained silenced for ~270 generations, resulting in cross progeny (F1) that were allowed to produce self-progeny of varying genotypes (F2) from which animals homozygous for *T* and for the wild-type or the mutant allele of *hrde-1* were assessed across generations by passaging self-progeny (F3 through F7). In addition, every generation of *hrde-1(-); T* hermaphrodites produced by self-fertilization (F2 through F6) was mated with either wild-type (+/+) or *hrde-1(-)* males to examine the possibility of re-initiation of transgenerational silencing. mCherry and GFP fluorescence was scored in heterozygous F1 cross progeny (*hrde-1(-/+)*) and in ≥F3 descendants of genotypes depicted. Cross progeny (grey text) of F2 *hrde-1(-); T* hermaphrodites mated with wild-type males were not obtained despite multiple biological repeats due to experimental design. Specifically, the mating was set up in replicates between a single *hrde-1(-); T* hermaphrodite with three wild-type males at every generation, beginning from the F2 generation onwards. The selection of hermaphrodites of *hrde-1(-); T* genotype was successful only from F3 generation, because homozygous *hrde-1(-); T* could only be set up from the F2 generation, which is the very first generation the genotype of descendants can become *hrde-1(-); T* after the cross set up at P0. As a result, because F2 *hrde-1(-); T* hermaphrodites were needed for crosses but *hrde-1(-); T* F2 animals could not be distinguished from their *hrde-1(+); T* or *hrde-1(+/-); T* siblings on the F1 > F2 plate. The only way to determine the genotype of the hermaphrodite used was by first mating a single random hermaphrodite of unknown *hrde-1* genotype with three wild-type males, and then allowing for the F3 progeny to be laid for 3 days before sacrificing the F2 hermaphrodite for genotyping. However, by this point, the F2 hermaphrodite, would be harbouring wild-type sperm in its spermatheca, confounding the genotyping PCR.

*Genetic crosses using animals overexpressing* gpr-1

To analyze DNA-independent signals we used a recently developed tool that prevents paternal and maternal pronuclei from fusing within the zygote[16,17]. A G protein regulator, GPR-1, when overexpressed maternally, increases forces that pull on spindle poles and prevents the maternal and paternal nuclei from fusing. This allows the contents of the paternal nucleus to be inherited into cells of the P lineage and the contents of the maternal nucleus to be inherited into the AB lineage. By way of such non-Mendelian segregation in most cross progeny, paternal DNA is inherited into all germline cells and select somatic cells (such as the intestine and body wall muscles) and maternal DNA is only inherited into the somatic cells (Fig. 2e). A smaller fraction of progeny either have maternal DNA in the germline and some soma and paternal DNA in most somatic cells (Fig. 2e) or undergo Mendelian segregation with paternal and maternal DNA in all cells (data not shown). To analyze the robustness of this tool in our hands, we tested the segregation of paternal and maternal DNA using *gtbp-1::gfp*, which expressed cytoplasmic GFP in all tissues (Fig. 2e). When hermaphrodites overexpressing *gpr-1* (*gpr-1 oe*) were crossed with males carrying *gtbp-1::gfp*, >95% of cross progeny showed non-Mendelian segregation with paternal DNA inherited into cells of the P lineage (based on presence of GFP in the germline) and showed segregation of maternal DNA into cells of the AB lineage (based on absence of GFP in some pharyngeal cells and neurons). A much smaller population of cross progeny (<5%) showed either the inverse pattern of segregation or Mendelian segregation. We used *gtbp-1::gfp* as the marker to identify non-mendelian cross progeny in further crosses with *gpr-1 oe*. To analyze effects of parental signals on *T* in the germline, we had to ensure that *T* (and the accompanying marker gene, *gtbp-1::gfp*) was always inherited from the male because the majority of non-Mendelian cross progeny would inherit paternal DNA into the germline. Since the transgene expressing *gpr-1* also expressed a synonymous variant of *gfp*, we used a variant of *T* i.e., $T\Delta\Delta\Delta$ or $Tcherry$ for further analyses to prevent GFP fluorescence from what would have been two different sources from confounding interpretation.

*Genetic crosses with Pmex-5::Tcherry::mex- 3' utr* and *Pmex-5::Tcherry::cye-1 3'utr*

Integration of *Pmex-5::Tcherry::mex-5 3' utr* and *Pmex-5::Tcherry::cye-1 3'utr* by MosSCI into the genome resulted in spontaneous silencing of the transgenes[18-20], whose expression could be revived by mutation of *hrde-1*. Because parental *hrde-1* was dispensable and zygotic *hrde-1* was sufficient for initiation of mating-induced silencing (Extended Data Fig. 6d), we used *Pmex-5::Tcherry::mex-5 3' utr; hrde-1(-)* or *Pmex-5::Tcherry::cye-1 3'utr; hrde-1(-)* parent animals in reciprocal crosses to test for mating-induced silencing (Fig. 4f), and scored cross progeny of genotypes *Pmex-5::Tcherry::mex-5 3' utr; hrde-1(+/-)* or *Pmex-5::Tcherry::cye-1 3'utr; hrde-1(+/-)*, respectively.

**Generation and maintenance of $iT$ and $iT\Delta$ strains**

To make hermaphrodites with $iT$ linked to a *dpy* marker, AMJ581 hermaphrodites were mated with N2 males to generate cross progeny males that all show bright mCherry fluorescence from *oxSi487*. These males were then mated with N2 hermaphrodites to give cross progeny (F1) with undetectable mCherry fluorescence. F1 animals were allowed to give progeny (F2) that were homozygous for *oxSi487* as determined by the homozygosity of a linked *dpy-2(e8)* mutation. One such F2 animal was isolated to be propagated as the *iT* strain (AMJ692).

To make males with *iT*, *dpy-17(e164) unc-32(e189)* hermaphrodites were mated with EG6787 males to generate cross progeny (F1) hermaphrodites with undetectable mCherry fluorescence. These cross progeny were allowed to give progeny (F2) that are homozygous for *oxSi487*. Two such F2s were isolated to be propagated as two different *iT* lines. One of these was designated as AMJ724 and used for further experiments. These strains maintained the silencing of *oxSi487* and were heat-shocked to produce males. Genotypes of *iT* strains were verified using PCR.

To make hermaphrodites with *iT*Δ linked to a *dpy* marker, AMJ767 hermaphrodites were mated with N2 males to generate cross progeny males with bright mCherry fluorescence. These males were then mated with GE1708 hermaphrodites to give cross progeny (F1) with undetectable mCherry fluorescence. F1 animals were allowed to give descendants that are homozygous for *T*Δ as determined by genotyping for *jamSi20*. A homozygous descendant was isolated to be propagated as the *iT*Δ strain (AMJ917). Genotypes of *iT*Δ strains were verified using PCR.

AMJ692 was used to test for recovery of gene expression ~150 generations after it was made. This generation time was estimated as follows: worms were passaged every 3.5 days for 143 generations over a period of 556 days, except for three intervals when they were allowed to starve and larvae were recovered after starvation. These intervals with recovery from starvation spanned a total of ~6 generations over 49 days. Thus, the total number of generations = 143 + ~6 = ~150 generations. The generation times for AMJ724, AMJ552 and AMJ844 were similarly estimated. *iT* strain silenced for >150 generations was used to test the requirements for RNAi factors in the maintenance of transgenerational silencing.

**CRISPR-Cas9 mediated editing of *oxSi487***

To generate edits in *oxSi487*, Cas9-based genome editing with a co-conversion strategy[51] was used. Guide RNAs were amplified from pYC13 using primers listed above. The amplified guides were purified (PCR Purification Kit, Qiagen) and tested in vitro for cutting efficiency (Cas9, New England Biolabs catalog no. M0386S). For most edits, homology template for repair (repair template) was made from gDNA using Phusion High Fidelity polymerase (New England Biolabs catalog no. M0530S) and gene specific primers to separately amplify regions precisely upstream and downstream of the site to be edited. The two PCR products were used as templates to generate the entire repair template using Phusion High Fidelity Polymerase and the fused product was purified using NucleoSpin Gel and PCR Clean-up (Macherey-Nagel, catalog no. 740609.250). Homology templates to generate *T*ΔΔ and *dpy-10(-)* were single-stranded DNA oligos. Wild-type animals were injected with 1.2 – 12.9 pmol/µl of guide RNAs, 0.08 – 1.53 pmol/µl of homology repair template to make edits in *T* and in *dpy-10* and 1.6 pmol/µl of Cas9 protein (PNA Bio catalog no. CP01). In animals with *T*ΔΔ edit, *Punc-119* deletion resulted in Unc animals due to the *unc-119(ed3)* mutation in the background of EG6787, suggesting that a functional transcript was not made from the remaining part of the rescuing *Punc-119::unc-119::unc-119 3'utr* insertion at *ttTi5605*. Edits were verified using PCR and Sanger sequencing. For additional details on specific reagents, see Extended Data Table 3.

**CRISPR-Cas9 mediated insertion**

To generate large insertions, the Cas9-based editing protocol was adapted from Dickinson *et al*, 2013[53]. The following mix was injected into HT1593 animals: 42-55 ng/µl plasmid expressing Cas9 protein and sgRNA sequence specific to chromosome II site near *ttTi5605* (pDD122) or chromosome I site near *ttTi4348* (pSD18), 105 ng/µl of pMA122 (*Phsp-16.41::peel-1::tbb-2utr*), 42-55 ng/µl of repair plasmid for insertion of $Tcherry^{Crispr}$ (*jamSi38, jamSi40, jamSi41*) or $Tcherry\ \mathrm{I}$ (*jamSi56*). Following injection, animals were singled out and the plate was allowed to crowd until starvation. Starved plates were heat shocked at 34°C for 2.5 to 4 hours and heat shocked animals were allowed to recover overnight. Non-Unc animals that survived the heat shock were singled out, propagated and screened for the edit using PCR. Single-copy insertions were then verified in isolates that screened positive for the edit after extraction of genomic DNA.

**Mos-mediated single copy insertion (MosSCI)**

To generate large insertions, the MosSCI protocol was adapted from Frøkjær-Jensen *et al*, 2012[7]. The following mix was injected into EG4322 animals: 50-55 ng/µl plasmid expressing Mos1 transposase (pCFJ601: *Peft-3::mos1 transposase::tbb-2utr*), 105 ng/µl of pMA122 (*Phsp-16.41::peel-1::tbb-2utr*), 50-

55 ng/µl of repair plasmid for insertion of *Tcherry*, *Tgfp*, *Tcherry-pi*, *Tcherry::tbb-2 3' utr* or *Tcherry::mex-5 3' utr* into chromosome II near *ttTi5605* insertion site. Following injection, animals were singled out and the plate was allowed to crowd until starvation. Starved plates were heat shocked at 34°C for 2.5 to 4 hours and heat shocked animals were allowed to recover overnight. Non-Unc animals that survived the heat shock were singled out, propagated and screened for the edit using PCR. Single-copy insertions were then verified in isolates that screened positive for the edit after extraction of genomic DNA.

**Quantitative RT-PCR (qPCR)**

Total RNA was isolated using TRIzol (Fisher Scientific) from 50-100µl pellets of mixed-stage animals. Three biological replicates were isolated by pelleting animals from three different plates of the same strain. RNA was extracted by chloroform extraction, precipitated using isopropanol, washed with ethanol and resuspended in 20-30 µl of nuclease-free water. 2-5 µl of resuspended RNA was set aside to run on a gel and the remaining was DNase-treated in DNase buffer (100 mM Tris-HCl, pH 8.5, mM $CaCl_2$, 25mM $MgCl_2$), and incubated with 0.25 µl DNase I (New England Biolabs, 2 units/µl) at 37°C for 60 minutes followed by heat inactivation and 75°C for 10 minutes. Pre- and post-DNase treated RNA were run on a 1% agarose gel to check for the presence of rRNA bands. RNA concentration was measured and equal amounts (500 ng to 1000 ng) of RNA were converted to cDNA using SuperScript III Reverse Transcriptase (Invitrogen catalog no. 18080044) with two-fold reduced quantities compared to manufacturer's recommendations. For cDNA conversion, 3-5 technical replicates were done for each biological replicate of each sample and RT primer P82 was used for *R11A8.1*, P83 for *tbb-2*, P84 for *mCherry* and P85 for *gfp*. qRT-PCR was done on cDNA using LightCycler 480 SYBR Green I Mastermix (Roche catalog no. 4707516001) guidelines according to the manufacturer's recommendations. For analysis of pre-mRNA, primers P86 and P87 were used for *R11A8.1*, P88 and P89 were used for *tbb-2*, P90 and P91 were used for *mCherry* and P92 and P93 were used for *gfp*. For analysis of mRNA, primers P94 and P95 were used for *tbb-2*, P96 and P97 were used for *mCherry* and P98 and P99 were used for *gfp*. Fold change was calculated using $2^{-Ct}$ values and samples were normalized to total RNA.

Three (Fig. 2h, Extended Data Fig. 6h) to six (Fig. 4e, Extended Data Fig. 10h) independent biological replicates were typically measured, with each biological replicate being the median of three to five technical replicates. A scaled scatter plot was used to depict the relative abundance of pre-mRNA and mRNA for each biological replicate. RNA abundance was estimated as proportional to $2^{-Cq}$ and target transcripts were normalized to total RNA to obtain relative abundance.

**Chromatin Immunoprecipitation-qPCR (ChIP-qPCR)**

This protocol was adapted from Guang *et al*[54]. 300 µl to 500 µl of frozen mixed-stage worm pellets were used for each ChIP experiment. Three biological replicates were done for every strain and worms from each sample were split into 100 µl pellets. Frozen pellets were crushed by grinding with a mortar and pestle. Crushed pellets were resuspended in 1 ml buffer A (15 mM Hepes-Na, pH 7.5, 60 mM KCl, 15 mM NaCl, 0.15 mM beta-mercaptoethanol (CALBIOCHEM catalog no. 444203), 0.15 mM spermine (Sigma-Aldrich catalog no. S3256-1G), 0.15 mM spermidine (Sigma-Aldrich catalog no. S2626-1G), 0.34M sucrose, 1XHALT protease (ThermoScientific catalog no. 78440) and phosphatase inhibitor cocktail (ThermoScientific catalog no. 78440)). To crosslink, formaldehyde was added to a final concentration of 2%, and incubated at room temperature for 15 minutes. The formaldehyde was quenched by adding 0.1 ml 1M Tris HCl (pH 8). The lysate was spun at 15,000g for 1 minute at 4°C. The resulting pellets were washed twice with ice-cold buffer A by centrifuging between washes. The pellets were resuspended in 0.3 ml buffer A with 2 mM $CaCl_2$. Micrococcal nuclease (Roche catalog no. M0247S ) was added to a final concentration of 0.3 U/µl and incubated for 5 minutes at 37°C (the tubes were inverted several times per minute). EGTA to a final concentration of 20 mM was added to stop the digestion reaction and samples were centrifuged at 15,000g for 1 minute at 4°C, followed by washing the resulting pellets with 300 µl of ice-cold RIPA buffer (1XPBS, 1% NP40 (Spectrum catalog no. T1279), 0.5% sodium deoxycholate (Sigma-Aldrich catalog no. D6750-10G), 0.1% SDS, 1XHALT protease and phosphatase inhibitor and 2 mM EGTA (Sigma-Aldrich catalog no. E3889-10G)). Samples were centrifuged at 15,000g for 1 minute at 4°C. The pellet was resuspended after washes in 0.8 ml ice-cold RIPA buffer, and solubilized by shearing using the Covaris[55]. Samples were kept on ice at all times except during shearing. All sheared lysates for each biological replicate were pooled and split equally to precipitate for all chromatin marks being measured. Sheared lysates were centrifuged at 15,000 g for 2 minutes. 80 µl of the supernatant was set aside at -20°C for "input" libraries and the remaining supernatant was used for IP. Antibodies were chosen based on their efficiency in *C. elegans*[56]. One of 2 µg of anti-H3 antibody (Abcam, ab1791), 3 µg of anti-H3K9me1 antibody (Abcam, ab8896), 3 µg of anti-H3K9me2 antibody (Abcam, ab1220) or 2 µg of anti-H3K9me3 antibody (Abcam, ab8898) was added and agitated gently at 4°C overnight. 50 µl of protein A Dynabeads (10% slurry in 1x PBS buffer) was added and mixed by shaking for 2 hours at 4°C. The beads were then washed four times (four minutes/wash) with ice-cold 600 µl LiCl washing buffer (100 mM Tris HCl, pH 8, 500 mM LiCl, 1% NP-40, 1% Sodium deoxycholate). A magnetic stand (DynaMag-2 Magnet, Thermo Scientific) was used to pellet beads and the supernatant was discarded after every wash. Beads and input were incubated with 450 µl worm lysis buffer (0.1 M Tris HCl, pH 8, 100 mM NaCl, 1% SDS) containing 200 µg/ml proteinase K at 65°C for 4 hours with agitation every 30 minutes to elute the immunoprecipitated nucleosome and reverse crosslinks. DNA was isolated by organic extraction and precipitation. DNA obtained was measured by qPCR (see qRT-PCR method) using LightCycler 480 SYBR Green I Mastermix according to the manufacturer's recommendations. Pre-mRNA primers (see qRT-PCR method) were used for analysis of *R11A8.1, mCherry* and *gfp*. Fold change was calculated using $2^{-\Delta\Delta Ct}$ method and samples were normalized to co-immunoprecipitated control gene, *R11A8.1*.

**Single molecule fluorescence *in situ* hybridization (smFISH)**

Custom Stellaris FISH probes were designed against only exons of *mCherry* and *gfp* sequence from *oxSi487* using the web-based Stellaris FISH Probe Designer from Biosearch Technologies (www.biosearchtech.com/stellarisdesigner). Any probe design expected to span exon-exon junctions was avoided to allow for the equivalent detection of both mature and nascent transcripts. Standard *C. elegans* smFISH protocol followed by 4',6-diamidino-2-phenylindole (DAPI) staining was used as described[57]. The probe blend to detect *mCherry* includes 25 exon-specific probes (P112 through P136) each tagged with Quasar 670 dye and antisense to *mCherry* RNA. The probe blend to detect *gfp* includes 26 exon-specific probes (P137 through P162) each tagged with Quasar 670 dye and antisense to *gfp* RNA. The adapted smFISH protocol is as follows: 50 to 100 L4 animals or adult animals ~24 hours post

L4 (Fig. 2c, Extended Data Fig. 7, Extended Data Fig. 10i) were paralyzed in 400 µl 1x Phosphate Buffered Saline 0.1% Tween-20 (PBST, Amresco, catalog number C999G23 K875-500ML) containing 0.25 mM levamisole for dissection or whole animals younger than L4 (Extended Data Fig. 7b) were washed in 1x PBST and fixed in 1 ml fix solution (3.7% formaldehyde (Amresco, catalog number 0493-500ML) in 1x PBST) on a nutator at room temperature. Fixation time ranged between 15 minutes and 45 minutes across different trials. Samples were washed in 1x PBST, incubated for 10 minutes. in permeabilizing solution (0.1% Triton X-100 in 1 ml of 1x Gibco PBS pH 7.4 (Thermofisher Scientific, catalog number 10010023)), washed twice in PBST and resuspended in 1 ml 70% ethanol and incubated between one to seven days at 4°C. Fixed animals were then equilibrated and washed with wash buffer (2x Sodium Saline Citrate (SSC, Sigma Aldrich, catalog number 11666681001), 10% formamide (Millipore Sigma, catalog number 4650-500ML or Amresco, catalog number 0314-500ML), 0.01% Tween-20 (Fisher Scientific, catalog number BP337-100)) hybridized with 0.025 µM probes diluted in hybridization buffer (10% dextran sulfate (Sigma Aldrich, catalog number D8906-5G), 2x SSC, 10% formamide) for 48 hours in a 37°C rotator in the dark. Hybridized animals were then washed in wash buffer, incubated with DAPI solution (1 µg/ml DAPI in wash buffer) for 30 minutes to 120 minutes. protected from light, washed twice in wash buffer for 5 minutes each in a rotator and used for mounting. Worms were resuspended and incubated for 5 minutes at room temperature or up to 6 hours at 4°C in a GLOX buffer without enzymes (2x SSC, 1% glucose (Fisher Scientific, catalog number D16-500), 0.1 M Tris pH 8.0 (Thermofisher Scientific, catalog number AM9855G) in RNase-free water), treated with freshly made GLOX-enzyme buffer (100 µl GLOX buffer, 1 µl glucose oxidase (MP Biomedicals/Fisher Scientific, catalog number 0219519610), 3.7 mg/ml, 1 µl catalase (Fisher Scientific, catalog number S25239A), 1 µl 200 mM Trolox (Acros Organics/Fisher Scientific, catalog number 218940050)) and prepared for imaging by dropping the sample on a coverslip followed by placing and sealing on a microscope slide with a mix of Vaseline, lanoline and paraffin. All samples within a single experimental set included control strains and were subjected to identical conditions (e.g. incubation times) to minimize variability within the experiment. RNase-free conditions were used in all smFISH experiments.

AMJ1259, AMJ1260 and AMJ1261 females were mated with AMJ1045 or EG6787 males and extruded gonads of cross progeny hermaphrodites staged at ~24 hours post L4 were subjected to smFISH protocol using *mCherry* probes (Extended Data Fig. 7c). For Extended Data Fig. 7d, e, extruded gonads of EG6787 ("*T*"), AMJ552 ("*iT*")) and N2 ("wild type") adult hermaphrodites staged at ~24 hours post L4 were subjected to the smFISH protocol using either *mCherry* or *gfp* probes. For Extended Data Fig. 10i top row, extruded gonads of EG6787, AMJ1170, JH3323 and N2 adult hermaphrodites staged at ~24 hours post L4 were subjected to the smFISH protocol using *mCherry* probes alone. For Extended Data Fig. 10i bottom row, extruded gonads of EG6787, AMJ1195, JH3197 and N2 adult hermaphrodites staged at ~24 hours post L4 were subjected to the smFISH protocol using *gfp* probes alone.

**Confocal microscopy to image single-molecule RNA signals or protein fluorescence**

Images were taken using Leica SP5 confocal microscope with the 63x oil immersion objective at 500% digital zoom for smFISH samples and 400% digital zoom to capture protein fluorescence. A single confocal slice of 0.5 µm thickness was captured at regions corresponding to distal, loop or proximal regions of the dissected gonad. The Z position was oriented to be the same plane as the nucleus of the distal tip cell for all three regions imaged in most dissected gonads. To image whole worms between L2 and L3 stages for smFISH, a Z stack of a part of the germline that could be accommodated within the field of view at the same magnification as was used for dissected gonads was imaged with a step size of 0.5 µm to 1 µm and displayed as a maximum intensity projection. Brightfield and DAPI images were taken using photomultiplier tubes whereas *mCherry* and *gfp* RNA and protein fluorescence images were taken using Hybrid Detector (HyD). For both smFISH and protein fluorescence, the XY laser scan was set to 400 Hz and imaged at a resolution of 1024 x 1024 pixels. Quasar 670 probes were excited using Alexa 633 nm laser (50% White Light Laser) and signal was detected between 650–715 nm with the pinhole at 105.05 µm. DAPI was excited using 405 nm (3-30% UV laser) and signal was acquired between 422–481 nm with the pinhole at 95.52 µm. For Quasar 670 and mCherry or GFP protein fluorescence, a line average of 6–8 with 1–2 frame accumulation was used. For DAPI, 3–4 line average was used.

**Quantification of smFISH signals**

Leica images (.lif format) were opened in Fiji (NIH), display range was adjusted, background was subtracted twice sequentially using a rolling ball radius of 50 pixels (~2.7 μm), threshold was adjusted, and number of RNA dots ≤ 250 object voxels in size were quantified per unit area. All parameters were adjusted identically among images of strains being compared. All images being compared were adjusted identically using Adobe Photoshop for display.

**Statistical analyses**

For each figure, $\chi^2$ test was used to compare data as indicated in figure legends except in cases where only one category (bright or silenced) was present in both datasets being compared. All comparisons shown include comparisons between only GFP fluorescence or only mCherry fluorescence within each experiment. Significance for ChIP and qRT-PCR experiments and crosses in Fig. 2a $Tgfp$, Extended Data Fig. 4e, Fig. 4 and Extended Data Fig. 6 were compared using Student's t-test.

**Genetic Inferences**

*Extent of mating-induced silencing is variable in progeny but is initiated in every mating.*

The initiation of mating-induced silencing is reliable (observed in >1500 animals from each one of >142 independent crosses in wild-type and *dpy*- or *unc*-marked genetic backgrounds). In every comparison, precisely the same markers were used in crosses being compared. Nevertheless, silencing (dim + off animals) varied from 68% to 100% in cross progeny in these backgrounds. The reason for this variation is unclear. Therefore, we did not strongly infer from small variations.

*Lack of silencing when the transgene is inherited only through self-sperm in hermaphrodites could be because of a protective signal transmitted through oocyte.*

Hemizygous self-progeny of hemizygous hermaphrodites showed stable expression of *T* for multiple generations (Extended Data Fig. 4c). In each generation the transgene is expected to be inherited through self-sperm 50% of the time and a maternal protective signal is required for expression of paternal *T* in genetic crosses (Fig. 3). Therefore, this result implies that either a protective signal inherited through oocytes licenses expression of *T* inherited through self-sperm in each generation or that inheritance of *T* through self-sperm does not result in silencing.

*The silencing signal can separate from $i$T in the male germline before meiotic maturation.*

While meiosis is completed in sperm before fertilization[58], it is stalled at prophase I in oocytes until fertilization[59]. Nevertheless, oocyte meiosis is completed early in the one-cell zygote such that only a haploid genome is present in the oocyte pronucleus when it meets the sperm pronucleus. Thus, a DNA-independent signal when transmitted through sperm must have separated from DNA in the male germline but when transmitted through oocytes can separate from DNA either in the hermaphrodite germline or in the embryo (Fig. 3d and Extended Data Fig. 10 b, c).

*Parental rescue of genes can complicate analysis of newly generated mutants*

Homozygous mutant progeny of heterozygous animals may not show the mutant defect because of rescue by parental gene products – typically maternal rescue. Consistently, only some *hrde-1(-/-)* progeny of *hrde-1(+/-)* animals showed expression but all *hrde-1(-/-)* progeny in the next generation showed expression (Extended Data Fig. 6f). All strains analyzed for initiation (Extended Data Fig. 6b) and maintenance (Extended Data Fig. 6e) requirements had been mutant for at least two generations, except when testing the requirement for *prg-1(-)* in initiation, which was done using *prg-1(-)* animals that were mutant for one generation.

**Supplemental Discussion**

**Comparison of mating-induced silencing with related epigenetic phenomena**

The hallmarks of mating-induced silencing are: (1) silencing is initiated upon inheritance only through the male sperm; (2) once initiated, silencing is stable for many generations; (3) transgenerational silencing is associated with a DNA-independent silencing signal that is made in every generation, can be inherited for one generation, and can silence homologous sequences; and (4) maternal exonic sequences can prevent initiation of silencing. While to our knowledge no other known phenomenon shares all of these hallmarks (Extended Data Table 2), phenomena that share some of these features are highlighted below and can inform future mechanistic studies.

Paramutation refers to meiotically heritable changes in gene expression transferred from one allele (“paramutagenic”) to another allele (“paramutable”) when they interact within a cell (reviewed in ref. 60). In addition to similar heritability, both paramutation[61-65] and mating-induced silencing rely on small RNAs to spread silencing from one locus to another homologous locus. However, there are several aspects of paramutation that were found to be different from mating-induced silencing, when tested. First, a paramutagenic allele often requires associated repetitive sequences[66-68]. Second, how a paramutagenic allele first arises remains obscure[60]. Third, while some alleles are paramutable, others are not, for reasons that are unknown[61]. The reliability of initiating and also protecting from meiotically heritable silencing at a defined single-copy locus described in this study will be useful in discovering possible shared mechanisms that have remained unclear in the ~60 years since the original discovery of paramutation in maize[62].

The unpredictable silencing that occurs at some single-copy reporter transgenes within the *C. elegans* germline has been called RNA-induced epigenetic silencing or RNAe[18,19,31,36,69]. Some studies of RNAe [18,69], but not others (p.94 in (19)) report genetic requirements for initiation and maintenance that are similar to those for mating-induced silencing – *prg-1* only for initiation and *hrde-1* for maintenance, although *hrde-1* was also required for initiation of mating-induced silencing. Transgenes silenced through RNAe are associated with specific genome sequences or a differential subset of small RNAs than are unsilenced transgenes[18,36,70] but it remains unclear whether these associated properties of the silenced loci are the cause or consequence of silencing. Nevertheless, a model proposing RNAe as a response to foreign or non-self DNA has emerged[18-20]. This model is inadequate because the same sequence can be either silenced or expressed within the germline (Fig. 1; ref. 18, 19, 36, 69) and endogenous genes are subjected to transgenerational silencing through similar PRG-1- and HRDE-1-dependent mechanisms[24,71-74]. Furthermore, the features of a transgene that trigger silencing are unknown. Tethering the Argonaute CSR-1 to the nascent transcript[35] or adding intronic sequences that are found in native germline-expressed genes[45] can increase the frequency of expression of a foreign sequence but does not itself determine whether a sequence is expressed. Thus, despite these efforts, the mechanisms that enable stable expression or silencing of a gene across generations remain unclear.

Unlike RNAe, mating-induced silencing can be predictably initiated and thus provides a reliable assay for evaluating how organisms establish stable expression or silencing of a gene. Our analyses suggest that the decision to express paternal foreign sequences (*mCherry* and *gfp*) is re-evaluated in each generation based upon maternal mRNA (Fig. 3). Although mating-induced silencing is not a general property of transgenes (Extended Data Fig. 3), a similar silencing phenomenon with dependence on maternal mRNA has been observed for the endogenous gene *fem-1* (ref. 27). However, it is unknown whether this *fem-1* silencing also shares the *trans* silencing properties and genetic requirements of mating-induced silencing.

Taken together, the paradigm of mating-induced silencing established here provides a reliable model to study epigenetic mechanisms that dictate expression or silencing of a sequence in every generation in otherwise wild-type animals.

**Implications for genetic studies**

The field of genetics relies heavily on analyses of animals generated by mating. Our study reveals that the direction of a genetic cross could strongly influence the phenotype of cross progeny. Additionally, because not every sibling from a cross has the same phenotype, the choice of the sibling selected for further manipulation can have a profound effect. Subsequent transgenerational persistence of silencing can make phenotype independent of genotype, resulting in erroneous conclusions. Thus, when using genetic crosses to generate strains both the direction of the genetic cross and choice of the individual cross progeny selected for propagation needs to be controlled for - especially when evaluating epigenetic phenomena. For example, we ensured that every cross was performed with the transgene present in the hermaphrodite to avoid initiating mating-induced silencing in our studies examining silencing by dsRNA from neurons[8]. Such methodological considerations impelled by this study could impact conclusions drawn from previous studies of epigenetic silencing in *C. elegans*.

**Possible impact on evolution**

Our results reveal a mechanism that silences genes in descendants in response to ancestral mating. The transgenerational stability of this gene silencing with the possibility of recovery of expression even after 170 generations (Fig. 2 and Extended Data Fig. 6) suggests that this mechanism could be important on an evolutionary time scale. Genes subject to such silencing could survive selection against their expression and yet be expressed in descendants as a result of either environmental changes that alter epigenetic silencing or mutations in the silencing machinery (e.g. in *hrde-1*). This mechanism thus buffers detrimental genes from selective pressures akin to how chaperones buffer defective proteins from selective pressures[75]. Many endogenous genes in *C. elegans* are silenced by HRDE-1 (ref. 18, 24, 74, 76), some of which could have been acquired when a male with the gene mated with a hermaphrodite without the gene. An interesting direction to explore next is to examine whether this mechanism facilitates adaptation.

**Data availability**

The data generated during and/or analysed during the current study are available from the corresponding author on reasonable request.

#### Extended Data Figures

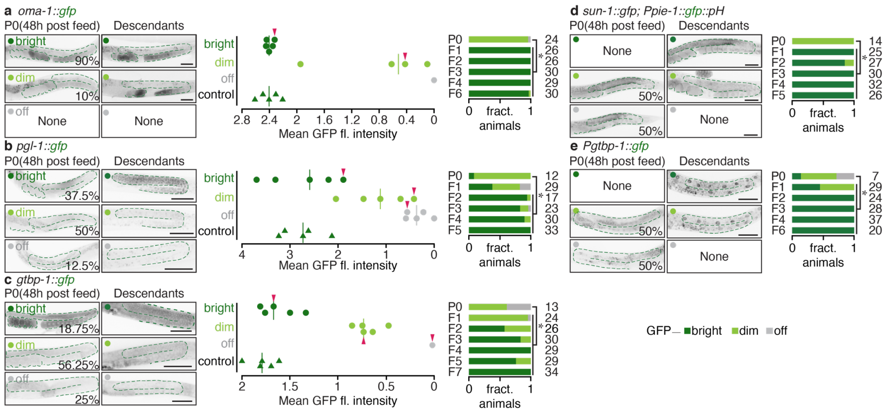


**Extended Data Figure 1. The same sequence can show variability in transgenerational silencing within the germline upon feeding RNAi.**

Five target genes expressing *gfp* (green) were exposed to control RNAi or dsRNA against *gfp* (*gfp* RNAi). The target genes were low copy (*Ppie-1::gfp::pH*, *oma-1::gfp*) or single copy (*Pmex-5::mCherry::gfp*) transgenes or endogenous gene tags (*gtbp-1::gfp*, *pgl-1::gfp*). Representative images of the germline (*far left*) of P0 animals exposed to RNAi for 24 hours and imaged an additional 24 hours later (48 hours post feed) to account for protein perdurance, are shown. Images of (*middle left*) and the level of GFP expression in (*middle right*) representative descendant animals (F1-F5) categorized as bright, dim or off are shown. Average (red line) normalized mean GFP fluorescence intensity within the germline was calculated for descendants of animals exposed to dsRNA against *gfp* (circles, bright: dark green, dim: light green, off: grey) or control dsRNA (green triangles). One to five L4-staged hermaphrodites were measured digitally after visually quantifying fluorescence from animals within each category. Red arrowheads indicate animals shown in representative images on the left. P0 animals (24 hours post feed) and F1-F5 descendants were analysed for expression of GFP and categorized based on intensity of fluorescence (*far right*) as in Fig. 2. The P0 to F7 data for *gtbp-1::gfp* (c) is the same as in Fig. 4d. Also see Fig. 1. Asterisks indicate $P < 0.05$ from $\chi^2$ test. Scale bar (50 µm) and number of animals scored (n) are indicated.

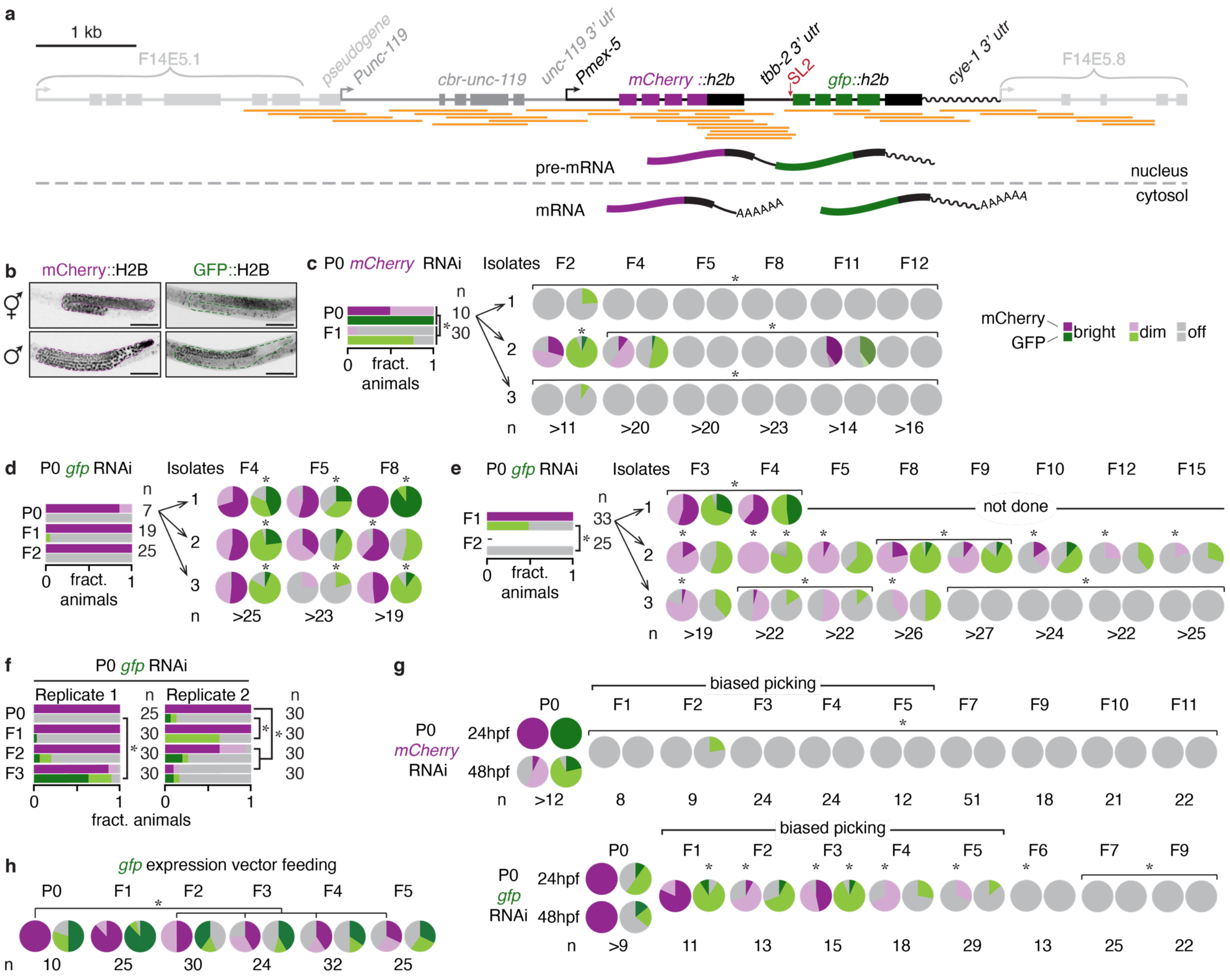

**Extended Data Figure 2. Dynamics of silencing of *mCherry* and *gfp* expressed from *T*.**

**a,** Schematic of *T* (*oxSi487*: *Pmex-5::mCherry::h2b::tbb-2 3' utr::gpd-2 operon::gfp::h2b::cye-1 3' utr*) within its genomic context where it is present as a single copy transgene as verified by PCR and Sanger sequencing. The transgene consists of *mCherry* and *gfp* genes tagged to *histone 2b* (*his-58*/*his-66*) arranged in an operon, and is presumably transcribed into one nascent transcript with both *mCherry::h2b* and *gfp::h2b* present as two separate mature transcripts in the cytosol. Orange lines correspond to stretches verified by individual Sanger sequencing experiments. The genes surrounding the insertion site of *T* on chromosome II are shown. **b**, Germlines (dotted outline) of representative L4-staged hermaphrodites and males showing mCherry::H2B or GFP::H2B expression from *T* are indicated. **c-g**, Animals expressing *T* were exposed to *mCherry* RNAi, *gfp* RNAi or control RNAi and scored for expression of mCherry and GFP for at least three generations. Early generations after P0 exposure to

RNAi were scored as in Extended Data Fig. 1. In (**c**-**e**) and (**g**), three animals were propagated in each generation and scored as explained in Fig. 1e, but in (**f**), twelve animals were propagated in every generation to reduce bottleneck effects and scored by imaging (see Methods). GFP expression was not scored in F2 animals in (**e**). Data in (**d**) and (**e**) is from animals exposed to the same RNAi food as those in Fig. 1e (*right*). Animals were blindly propagated in every generation (**c-f**) or silenced animals scored by eye were propagated (biased picking) for up to five generations and then blindly propagated in subsequent generations (**g**). In **g**, animals imaged an additional 24 hours post feeding RNAi (48 hpf) showed further decrease in mCherry or GFP expression suggestive of protein perdurance 24 hours post feeding RNAi (24 hpf). **h**, Animals expressing *T* were exposed to bacteria carrying a *gfp* expression vector or control RNAi and scored for expression of mCherry and GFP for five generations. Animals were propagated in an unbiased manner. In all figures, P0 animals exposed to control RNAi and their descendants showed bright expression of mCherry and GFP. Also see Fig. 1. Number of animals assayed and scale bar are as in Fig. 1. Asterisks are as in Extended Data Fig. 1 and indicate significant differences upon comparison to P0 animals (**c-f**, **h**) or 48 hpf P0 animals (**g**).

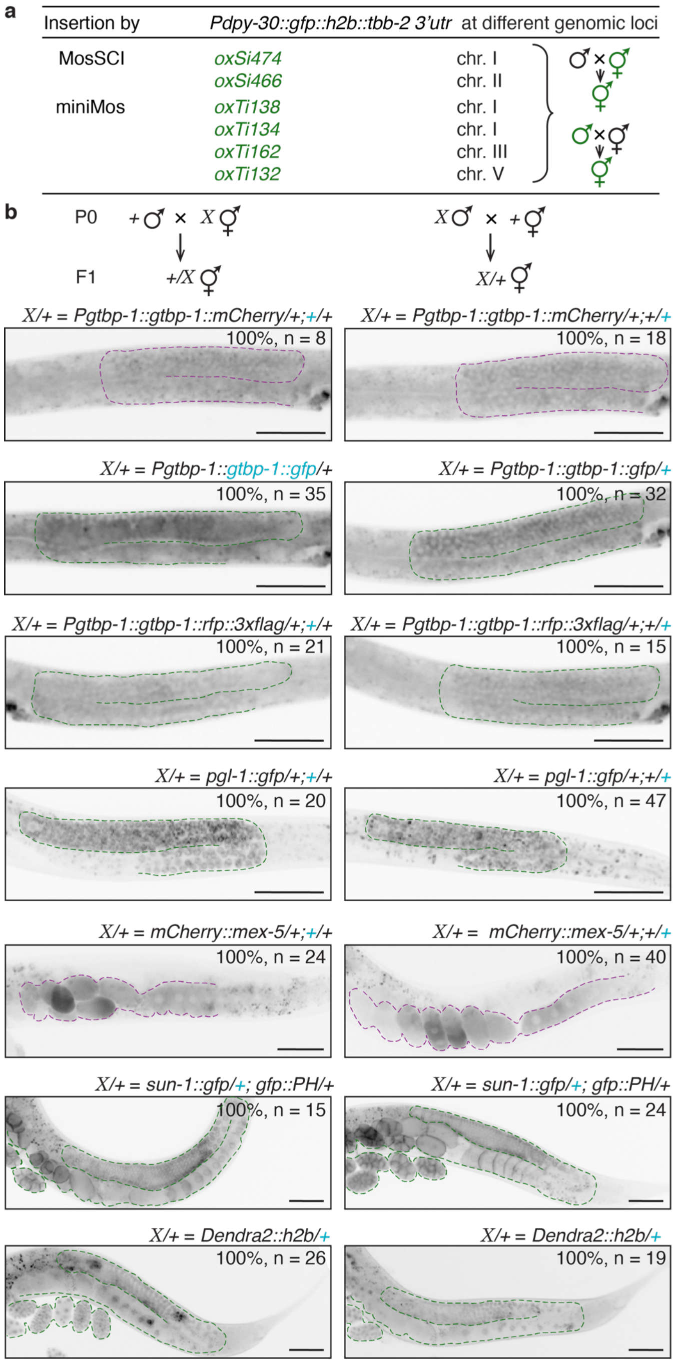


**Extended Data Figure 3. Expression within the germline remains unaffected by mating for many tested genes.**

Transgenes made using miniMos[45] (*Pdpy-30::gfp::h2b::tbb-2 3' utr*), MosSCI (*Pdpy-30::gfp::h2b::tbb-2 3' utr*, *sun-1::gfp* and *Pmex-5::Dendra2::h2b::tbb-2 3' utr*), or bombardment (*Ppie-1::gfp::PH(PLCdelta1)*) and endogenous genes tagged with reporter sequences using CRISPR-Cas9-mediated genome editing (*gtbp-1::gfp*, *mCherry::mex-5*, *gtbp-1::rfp::3xflag*, *pgl-1::gfp*, and *gtbp-1::mCherry*), or bombardment

(*Ppie-1::gfp::PH(PLCdelta1)*) were tested for susceptibility to mating-induced silencing as in Fig. 2a. Germlines of representative cross progeny at L4 or adult stage are outlined in **b**. Number of animals assayed, scale bar and blue font are as in Fig. 2.

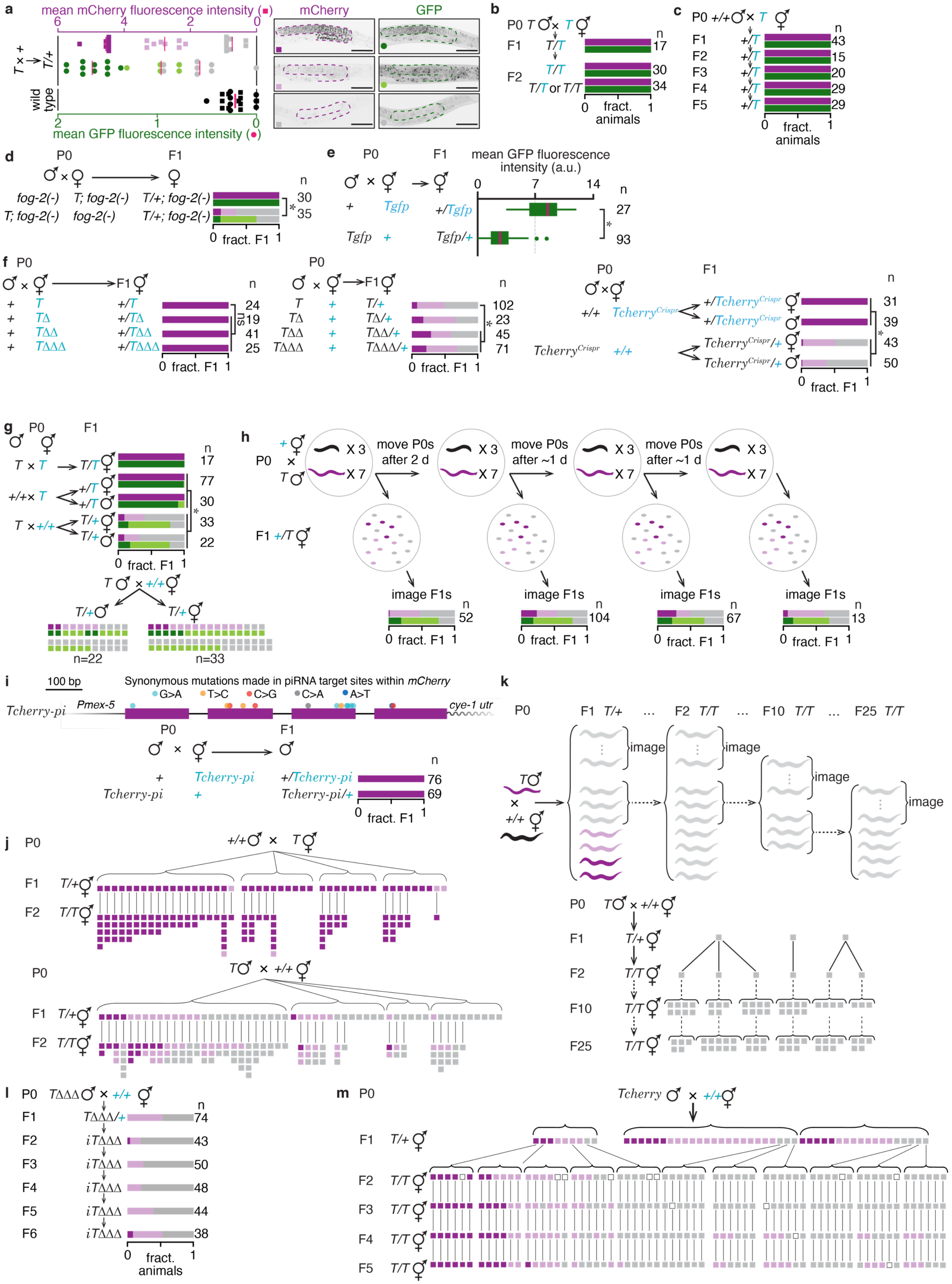
a
mean mCherry fluorescence intensity
6 4 2 0
T × +
T/+
wild type
2 1 0
mean GFP fluorescence intensity
mCherry
GFP
b
P0 T × T
F1 T/T
F2 T/T
T/T or T/T
n
17
30
34
0 fract. animals 1
c
P0 +/+ × T
F1 +/T 43
F2 +/T 15
F3 +/T 20
F4 +/T 29
F5 +/T 29
n
0 fract. animals 1
d
P0 F1
fog-2(-) T; fog-2(-) T/+; fog-2(-)
T; fog-2(-) fog-2(-) T/+; fog-2(-)
n
30
35
*
0 fract. F1 1
e
P0 F1
mean GFP fluorescence intensity (a.u.)
0 7 14
+ Tgfp +/Tgfp
Tgfp + Tgfp/+
n
27
93
*
f P0 F1
+ T +/T 24
+ TΔ +/TΔ 19
+ TΔΔ +/TΔΔ 41
+ TΔΔΔ +/TΔΔΔ 25
ns
0 fract. F1 1
P0 F1
T + T/+ 102
TΔ + TΔ/+ 23
TΔΔ + TΔΔ/+ 45
TΔΔΔ + TΔΔΔ/+ 71
*
0 fract. F1 1
P0 F1
+/+ Tcherry^Crispr
+/Tcherry^Crispr 31
+/Tcherry^Crispr 39
Tcherry^Crispr +/+
Tcherry^Crispr/+ 43
Tcherry^Crispr/+ 50
*
0 fract. F1 1
g P0 F1
T × T T/T 17
+/+× T +/T 77
+/T 30
T × +/+ T/+ 33
T/+ 22
*
n
0 fract. F1 1
T × +/+
T/+
T/+
n=22
n=33
h
+
P0
T
X 3
X 7
move P0s after 2 d
move P0s after ~1 d
move P0s after ~1 d
F1 +/T
image F1s
n
52
104
67
13
0 fract. F1 1
i
100 bp
Synonymous mutations made in piRNA target sites within mCherry
G>A T>C C>G C>A A>T
Tcherry-pi
Pmex-5
cye-1 utr
P0 F1
+ Tcherry-pi +/Tcherry-pi 76
Tcherry-pi + Tcherry-pi/+ 69
n
0 fract. F1 1
j P0 +/+ × T
F1 T/+
F2 T/T
P0 T × +/+
F1 T/+
F2 T/T
k
P0 F1 T/+ ... F2 T/T ... F10 T/T ... F25 T/T
T
×
+/+
image
P0 T × +/+
F1 T/+
F2 T/T
F10 T/T
F25 T/T
l P0 TΔΔΔ × +/+
F1 TΔΔΔ/+ 74
F2 iTΔΔΔ 43
F3 iTΔΔΔ 50
F4 iTΔΔΔ 48
F5 iTΔΔΔ 44
F6 iTΔΔΔ 38
n
0 fract. animals 1
m P0 Tcherry × +/+
F1 T/+
F2 T/T
F3 T/T
F4 T/T
F5 T/T

**Extended Data Figure 4. Mating-induced silencing is piRNA-dependent and results in transgenerational epigenetic inheritance.**

**a,** Quantification (*left*) and representative images (*right*) of the germline (magenta outline) of hemizygous animals (*T/+*) scored as having bright (*top*), dim (*middle*), or not detectable (off, *bottom*) levels of mCherry or GFP fluorescence. Average (red bar) normalized fluorescence within the germline was calculated for 11 bright, 5 to 8 dim, 8 off (grey), and 7 wild-type (black) L4-staged hermaphrodites. **b**, Males and hermaphrodites expressing *T* were mated, and fluorescence was scored in cross progeny (F1) and self-fertilized grand-progeny (F2) that inherited only the grand-maternal allele or only the grand-paternal allele or both. F1 data shown here is the same as that in (**g**). **c**, Wild-type males were mated with *T* hermaphrodites and hemizygous cross progeny (F1) as well as in descendant hemizygous self-progeny (F2 through F5) were scored. In contrast to previous reports[77], we find that *T* is not subject to meiotic silencing by unpaired DNA[78]. **d**, Mutation of *fog-2* feminizes the germline in 100% of hermaphrodites but has no effect in males. Feminized mothers were used in a control cross or in a cross to initiate mating-induced silencing. **e**, Germline GFP fluorescence from hemizygous *Tgfp*/+ cross progeny from Fig. 2a was quantified. **f**, Animals expressing variants of *T* were mated with non-transgenic animals and cross progeny were scored. **g**, Cross progeny males and hermaphrodites that inherited *T* from one or both parents were scored. Scoring data from the cross is re-plotted below to show mCherry and GFP fluorescence in each individual (colored box pair). **h**, *T* males and non-transgenic hermaphrodites were mated and cross progeny that were laid in the first 48 hours (2 days, 2 d) or in subsequent ~24 hours (1 day, 1 d) intervals, were collected after moving the P0s at these intervals to fresh plates. While silencing triggered by parental ingestion of dsRNA is less effective in later progeny[5,6], silencing triggered by mating can be equally effective in early and in late progeny. **i**, Schematic of synonymous changes in predicted piRNA sites within *mCherry* is depicted. Animals expressing *Tcherry* without piRNA binding sites (*Tcherry-pi*) were mated with non-transgenic animals, and cross progeny males were scored. **j**, Animals expressing *T* were mated with wild-type animals in four independent crosses (brackets) and mCherry fluorescence was scored in hemizygous cross progeny and in homozygous grand-progeny. Each box indicates fluorescence intensity (as in **a**) of a single adult animal and lines indicate descent. Once initiated, mating-induced silencing persists despite passage of *T* through oocytes of hermaphrodites and is therefore unlike genomic imprinting[79,38], where passage of *T* through oocytes is expected to revive expression. **k**, F2 ‘off’ progeny (from **j**) obtained after initiation of mating-induced silencing were propagated without further selection by selfing for 23 generations as indicated by the passaging scheme. mCherry fluorescence intensity was measured in animals (boxes) at F1, F2, F10 and F25 generations from three independent P0 crosses. At each generation indicated, siblings of the animals that were passaged were scored. Presence of the transgene was verified by genotyping in F1 and F2 generations. **l**, *T*ΔΔΔ males were mated with non-transgenic hermaphrodites and scoring was done in cross progeny (F1) and in descendants propagated blindly from ‘off’ F1 animals. **m**, *Tcherry* males were mated with non-transgenic hermaphrodites in three independent crosses and cross progeny belonging to each fluorescence level were singled out to give F2 animals. From F2 through F5, a single animal was blindly passaged and a single descendant was scored. Empty box indicates that the animal could not be scored because it was lost after being passaged on to a fresh plate, but only after having laid eggs, which enabled the continued scoring of its descendants. In all panels, scoring of silencing, number of animals assayed, scale bars and blue font are as in Fig. 2a. ‘ns’, statistically not significant. Asterisks indicate $P < 0.05$ from $\chi^2$ test (**d**, **f**, **g**) or Student’s t-test (**e**).

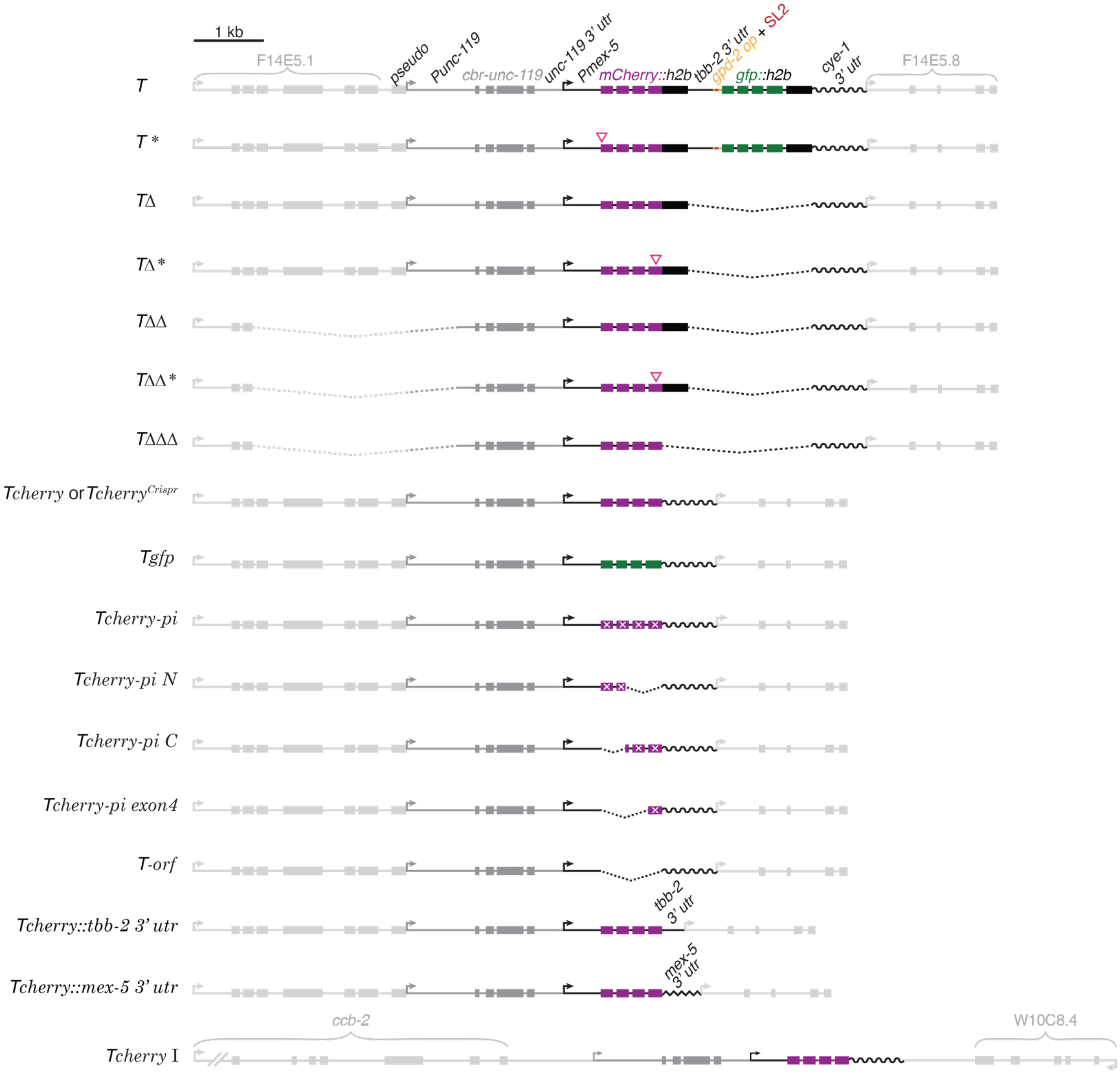


**Extended Data Figure 5. Schematics of *T*, of serial deletions and/or indels of *T* and of minimal variants of *T* that were newly integrated into a naive genome.** Schematic of *Pmex-5::mCherry::h2b::tbb-2 3'utr::gpd-2 operon::gfp::h2b::cye-1 3' utr* transgene (called *T* in this study). Successive deletions that remove *gfp* and *tbb-2 3' utr* (*T*Δ), a ~3 kb region upstream of the *unc-119(+)* coding region (*T*ΔΔ), and *h2b* (*T*ΔΔΔ) are depicted in their genomic context, along with variations that in addition contain small indels (*T**, *T*Δ*, *T*ΔΔ*). *Tcherry*, *Tcherry*$^{Crispr}$, *Tgfp*, *Tcherry::tbb-2 3' utr*, *Tcherry::mex-5 3' utr* and *Tcherry* on chromosome I were integrated independently of each other.

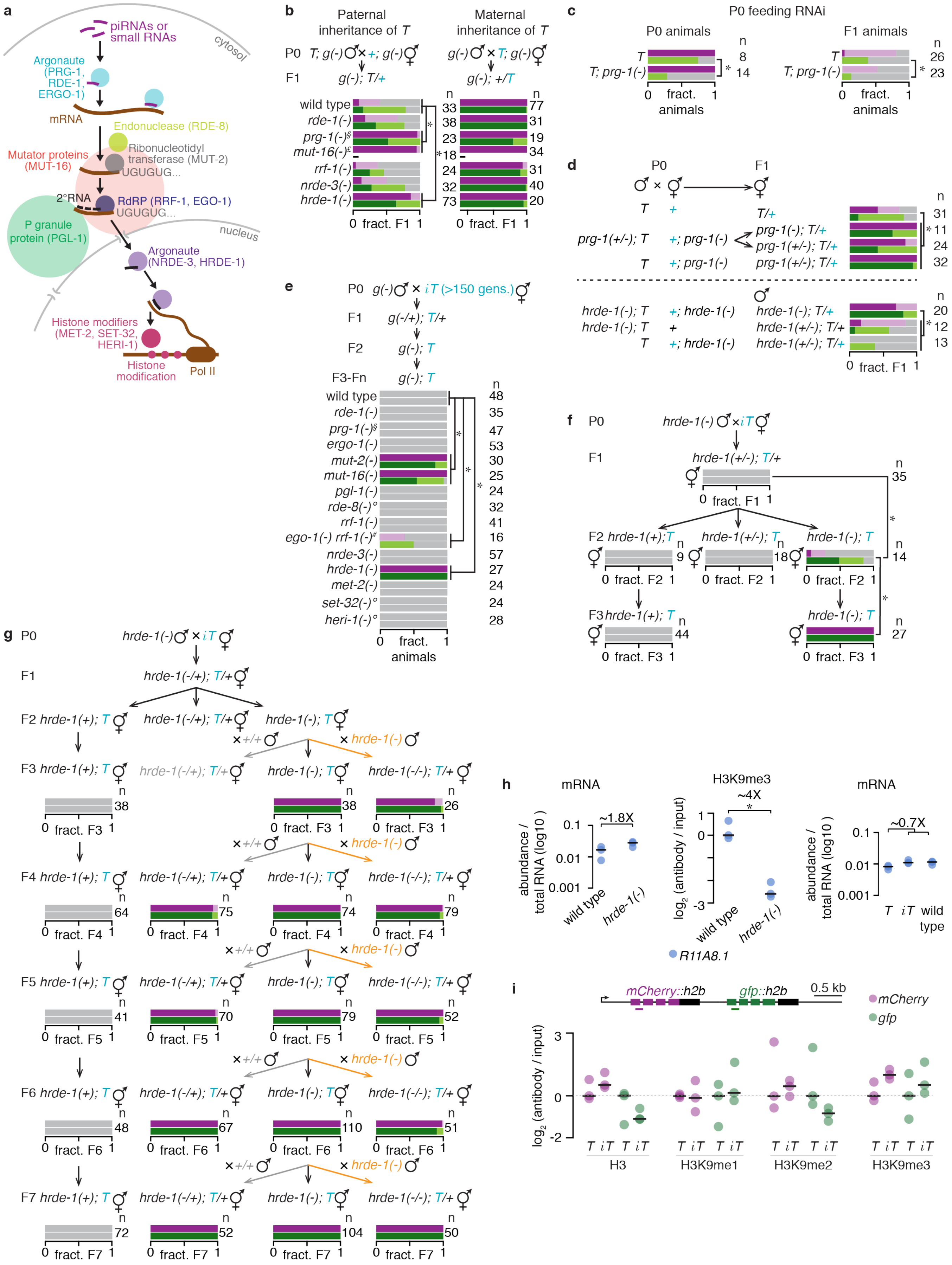

a
piRNAs or small RNAs
cytosol
Argonaute (PRG-1, RDE-1, ERGO-1)
mRNA
Endonuclease (RDE-8)
Mutator proteins (MUT-16)
Ribonucleotidyl transferase (MUT-2)
UGUGUG...
2°RNA
RdRP (RRF-1, EGO-1)
UGUGUG...
P granule protein (PGL-1)
nucleus
Argonaute (NRDE-3, HRDE-1)
Histone modifiers (MET-2, SET-32, HERI-1)
Histone modification
Pol II
b
Paternal inheritance of T
Maternal inheritance of T
P0
F1
wild type
rde-1(-)
prg-1(-)§
mut-16(-)£
rrf-1(-)
nrde-3(-)
hrde-1(-)
n
33 38 23 18 24 32 73
77 31 19 34 31 40 20
0 fract. F1 1
c
P0 feeding RNAi
P0 animals
F1 animals
T
T; prg-1(-)
n
8
14
26
23
0 fract. animals 1
d
P0
F1
T/+
prg-1(-); T/+
prg-1(+/-); T/+
prg-1(+/-); T/+
n
31 11 24 32
hrde-1(-); T/+
hrde-1(+/-); T/+
hrde-1(+/-); T/+
n
20 12 13
0 fract. F1 1
e
P0 g(-) × iT (>150 gens.)
F1 g(-/+); T/+
F2 g(-); T
F3-Fn g(-); T
wild type
rde-1(-)
prg-1(-)§
ergo-1(-)
mut-2(-)
mut-16(-)
pgl-1(-)
rde-8(-)°
rrf-1(-)
ego-1(-) rrf-1(-)#
nrde-3(-)
hrde-1(-)
met-2(-)
set-32(-)°
heri-1(-)°
n
48 35 47 53 30 25 24 32 41 16 57 27 24 24 28
0 fract. animals 1
f
P0 hrde-1(-) × iT
F1 hrde-1(+/-); T/+
n 35
F2 hrde-1(+); T
hrde-1(+/-); T
hrde-1(-); T
n 9 18 14
F3 hrde-1(+); T
hrde-1(-); T
n 44 27
0 fract. F3 1
g
P0 hrde-1(-) × iT
F1 hrde-1(-/+); T/+
F2 hrde-1(+); T
hrde-1(-/+); T/+
hrde-1(-); T
×+/+
× hrde-1(-)
F3 hrde-1(+); T
hrde-1(-/+); T/+
hrde-1(-); T
hrde-1(-/-); T/+
n 38 38 26
F4 n 64 75 74 79
F5 n 41 70 79 52
F6 n 48 67 110 51
F7 n 72 52 104 50
h
mRNA
abundance / total RNA (log10 )
~1.8X
wild type
hrde-1(-)
H3K9me3
~4X
log2 (antibody / input)
R11A8.1
mRNA
~0.7X
T iT wild type
i
mCherry::h2b
gfp::h2b
0.5 kb
mCherry
gfp
log2 (antibody / input)
T iT T iT
H3
H3K9me1
H3K9me2
H3K9me3

**Extended Data Figure 6. Genetic requirements for initiation and maintenance of mating-induced silencing.**

**a**, Schematic depicting the described role of different components of the RNAi pathway that were examined for their requirement in initiation or maintenance of mating-induced silencing[12-14]. Within the germline, 2º RNA production can be uncorrelated with gene silencing[11,23]. **b**, Mating-induced silencing was initiated as in Fig. 2a in a wild-type or in different mutant (*g(-)*) backgrounds (*left*) and silencing in resulting cross progeny were compared with that of the same genotypes from control crosses (*right*). Asterisk indicates $P < 0.05$ for a comparison with cross done in the wild-type background. Wild-type crosses shown here are the same as in Extended Data Fig. 4g. An additional wild-type cross with a different visible marker (mCherry: bright = 5, dim = 6, off = 25 and GFP: bright = 7, dim = 12, off = 17) was performed for comparison with the *rde-1(-)* cross on the right. Requirement of *mut-16* in initiation of silencing was examined by scoring only mCherry fluorescence in male cross progeny (£, see Methods). **c**, Animals expressing *T* in a wild-type or *prg-1(-)* background were exposed to *gfp* RNAi or control RNAi for one generation as in Fig. 1a and their untreated progeny were scored. **d**, Requirement of *prg-1* and *hrde-1* in initiation was tested by mating parents mutant for either of these genes and scoring cross progeny. **e**, $iT$ hermaphrodites after 150 to 250 generations of silencing were mated with males mutant for RNAi components (*g(-)*) and resulting descendants homozygous for the mutant allele of the gene were scored. Use of *prg-1(-/+)* males (§) owing to the poor mating by *prg-1(-)* males in (**b**) and (**f**) is indicated. Use of fertile *ego-1(-/+) rrf-1(-/+)* hermaphrodites, rather than sterile *ego-1(-) rrf-1(-)* hermaphrodites and $iT$ males (#) is indicated. **f**, *hrde-1(-)* mutants were mated with $iT$ silenced for 171 generations, and scoring was performed in cross progeny, in F2 and F3 descendants. **g**, Experiment depicting the test for whether $iT$ that recovers expression upon removal of *hrde-1(-)* (orange) can show silencing upon re-introduction of *hrde-1(+)* (grey) without re-initiating mating-induced silencing in the descending generations. F3 animals of the genotype *hrde-1(+/-); T/+* from F2 *hrde-1(-); T* hermaphrodites crossed with N2 males were not obtained due to experimental constraints. **h**, RT-qPCR of mRNA and ChIP-qPCR of H3K9me3 levels of an *hrde-1* target gene[18,24], *R11A8.1*, were measured in wild-type, *hrde-1(-)*, *T* and $iT$ animals. H3K9me3 measurements were normalized to wild-type levels. Similar to previous reports, we detected a decrease in H3K9me3 at the *R11A8.1* gene upon loss of HRDE-1, however, no significant change in mRNA was detected. mRNA levels of *R11A8.1* was not significantly altered between *T*, $iT$ and wild-type animals and hence was used as a control gene for ChIP experiments. Each filled dot represents one biological replicate and black line indicates the median value. Each mRNA measurement is the median of five technical replicates. **i**, H3, H3K9me1, H3K9me2 and H3K9me3 levels were measured at genomic *mCherry* and *gfp* in *T* and $iT$ animals. Measurements were normalized to levels at *R11A8.1* measured from each sample's respective input and then to *T*. Each filled circle represents one biological replicate, which is the median of five technical replicates and black line indicates the median value. In all panels, scoring of silencing, number of animals assayed, and blue font are as in Fig. 2a. Asterisks indicate $P < 0.05$ from $\chi^2$ test, Wilson's estimates for proportions (**e**) or Student's t-test (**h, i**).

Also see 'Genetic Crosses' under Methods.

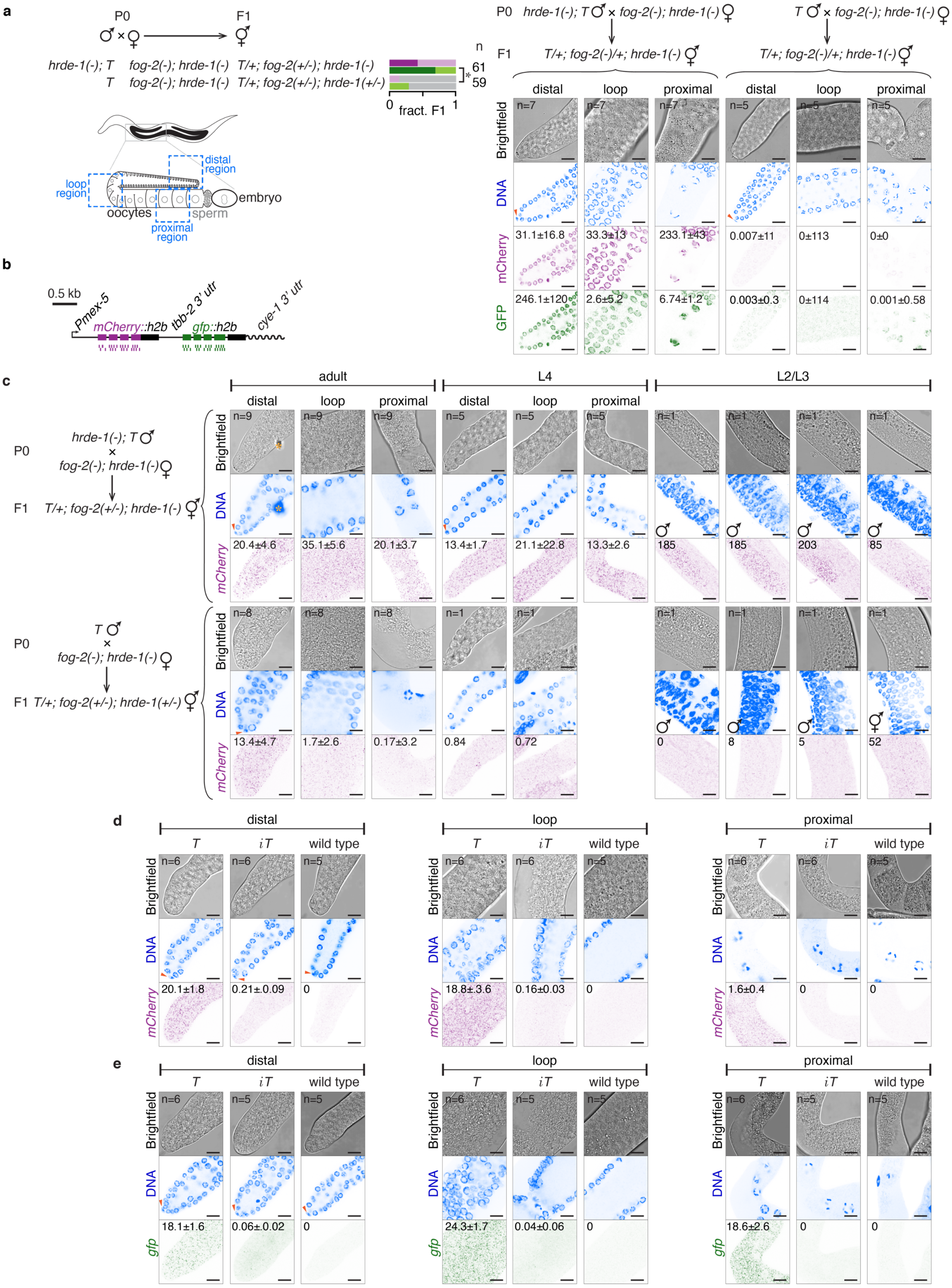
a
P0
F1
hrde-1(-); T
fog-2(-); hrde-1(-)
T/+; fog-2(+/-); hrde-1(-)
T
fog-2(-); hrde-1(-)
T/+; fog-2(+/-); hrde-1(+/-)
n
61
59
*
0
fract. F1
1
distal region
loop region
oocytes
sperm
embryo
proximal region
hrde-1(-); T ♂ × fog-2(-); hrde-1(-) ♀
T/+; fog-2(-)/+; hrde-1(-) ⚥
T ♂ × fog-2(-); hrde-1(-) ♀
T/+; fog-2(-)/+; hrde-1(-) ⚥
distal
loop
proximal
Brightfield
DNA
mCherry
GFP
n=7
n=5
31.1±16.8
33.3±13
233.1±43
0.007±11
0±113
0±0
246.1±120
2.6±5.2
6.74±1.2
0.003±0.3
0±114
0.001±0.58
b
0.5 kb
Pmex-5
mCherry::h2b
tbb-2 3' utr
gfp::h2b
cye-1 3' utr
c
adult
L4
L2/L3
distal
loop
proximal
P0
hrde-1(-); T ♂
×
fog-2(-); hrde-1(-) ♀
F1 T/+; fog-2(+/-); hrde-1(-) ⚥
n=9
n=5
n=1
20.4±4.6
35.1±5.6
20.1±3.7
13.4±1.7
21.1±22.8
13.3±2.6
185
185
203
85
T ♂
×
fog-2(-); hrde-1(-) ♀
F1 T/+; fog-2(+/-); hrde-1(+/-) ⚥
n=8
n=1
13.4±4.7
1.7±2.6
0.17±3.2
0.84
0.72
0
8
5
52
d
distal
loop
proximal
T
iT
wild type
n=6
n=5
20.1±1.8
0.21±0.09
0
18.8±3.6
0.16±0.03
0
1.6±0.4
0
0
e
gfp
18.1±1.6
0.06±0.02
0
24.3±1.7
0.04±0.06
0
18.6±2.6
0
0

**Extended Data Figure 7. Mating-induced silencing occurs by quantitative reduction of both *mCherry* and *gfp* transcripts and protein within the germline in cross progeny and across generations.** **a**, *T* or *T; hrde-1(-)* males were mated with *hrde-1(-); fog-2(-)* females and fluorescence due to mCherry::H2B and GFP::H2B in cross progeny was scored (*left top*) by eye or using confocal slices of indicated regions of dissected gonads (*right*). Scoring of silencing and number of animals assayed are as in Fig. 2a. Schematics of imaged regions (**a**) and single-molecule fluorescence *in situ* hybridization (smFISH) probes that hybridize to *mCherry* or *gfp* exonic RNA (**b**) are indicated. **c**, smFISH of *mCherry* in cross progeny adults obtained from a mating as in (**a**). Images of distal region in adults are also shown in Fig. 2c. **d-e**, smFISH of *mCherry* (**d**) or *gfp* (**e**) exonic RNA was performed in indicated regions of dissected gonads of adult wild-type, *T* or $iT$ animals. Pink arrowheads indicate the nucleus of the distal tip cell (**a-e**) and orange asterisks indicate non-specific signal (**c-e**). Numbers within images refer to mean fluorescence intensity per unit area measured in arbitrary units (**a**) or number of RNAs per 100 $\mu m^2$ (**c-e**) with standard error of the mean. Animals with median values of fluorescence or RNA signal in the distal region are shown in representative images along with the loop and proximal regions (**a, c-e**) within the same animals. Scale bar, 8 μm (**a**) or 10 μm (**c-e**). Number of animals imaged per region is indicated within the brightfield image.

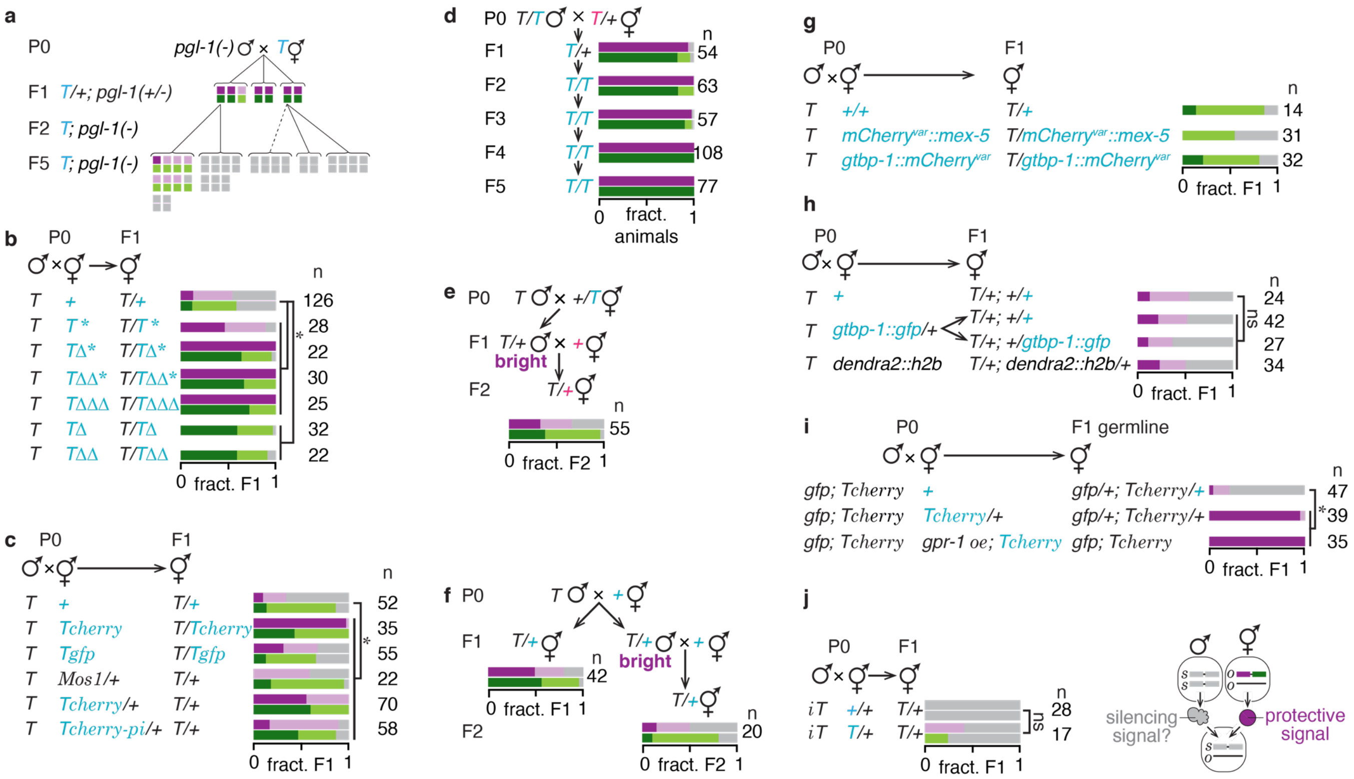

**Extended Data Figure 8. Maternal signals from *T* can prevent mating-induced silencing but cannot robustly reverse transgenerational silencing.**

**a**, *T* animals were mated with *pgl-1* mutants and expression of *T* was assessed in hemizygous cross progeny and in homozygous descendants. **b**, **c**, *T* males were mated with hermaphrodites containing a variant of *T* and paternally inherited *T* in resulting cross progeny males was scored. **d**, *T* hermaphrodites were mated with wild-type males and hemizygous cross progeny (F1) as well as four generations of homozygous descendants (F2 through F5) were scored. **e-f**, Male progeny with bright mCherry fluorescence that were protected from initiation (**e**) or that escaped initiation of mating-induced silencing (**f**) were subjected to mating-induced silencing. **g-h**, Males expressing *T* were mated with hermaphrodites expressing genes with homologous protein (**g**) or DNA (**h**) sequences, and fluorescence from paternally inherited *T* was scored in cross progeny. **i**, Males expressing *Tcherry*; *gtbp-1::gfp* were mated with hermaphrodites that expressed *Tcherry* in a wild-type or *gpr-1* overexpression (*oe*) background and fluorescence of paternally inherited *Tcherry* was scored in cross progeny. **j**, *iT* males were mated with non-transgenic or hemizygous hermaphrodites and cross progeny inheriting only paternal *iT* were scored. Scoring of silencing, number of animals assayed, and blue or pink font are as in Fig. 3a. Asterisks indicate $P < 0.05$ from $\chi^2$ test. 'ns' indicates no significant difference using $\chi^2$ test.

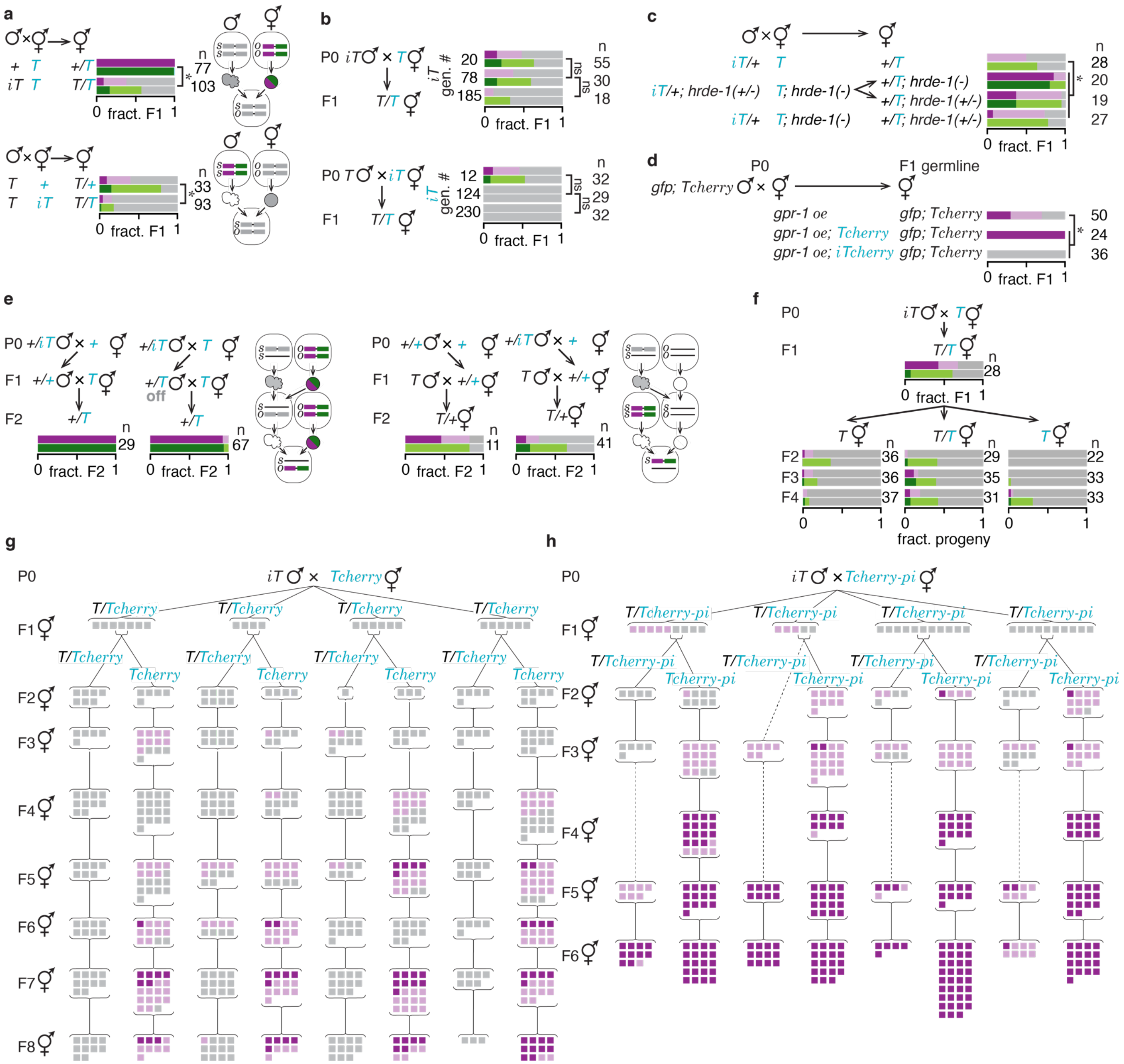


**Extended Data Figure 9. Recovery of expression from silencing *in trans* is locus-specific.**

**a, b**, Animals expressing *T* were mated with *iT* animals that remained silenced for many generations (*iT* gen. #), and cross progeny were scored. The combined data from each cross in (**b**) is shown in (**a**). **c**, Requirement of *hrde-1* for the activity of the silencing signal was tested by parental or maternal removal of HRDE-1. **d**, *Tcherry* animals silenced for more than five generations upon mating-induced silencing were designated as *iTcherry*. Males expressing *Tcherry; gtbp-1::gfp* were mated with hermaphrodites with *gpr-1* overexpression with or without *iT cherry* or *Tcherry*. Expression of paternally inherited *Tcherry* in the germline was scored in cross progeny. **e**, Crosses to test the transmission of the separable silencing signal across more than one generation. **f-h**, *T*, *Tcherry* or *Tcherry-pi* were mated with *iT* animals and resulting cross progeny and subsequent generations of descendants were scored for maternally inherited mCherry. GFP fluorescence was off in all scored animals (data not shown), independent of the level of fluorescence of mCherry fluorescence from *Tcherry* or *Tcherry-pi*. In all panels, scoring of silencing, number of animals assayed, and blue font are as in Fig. 2a. Asterisks indicate $P < 0.05$ and 'ns' indicates no significant difference from $\chi^2$ test.

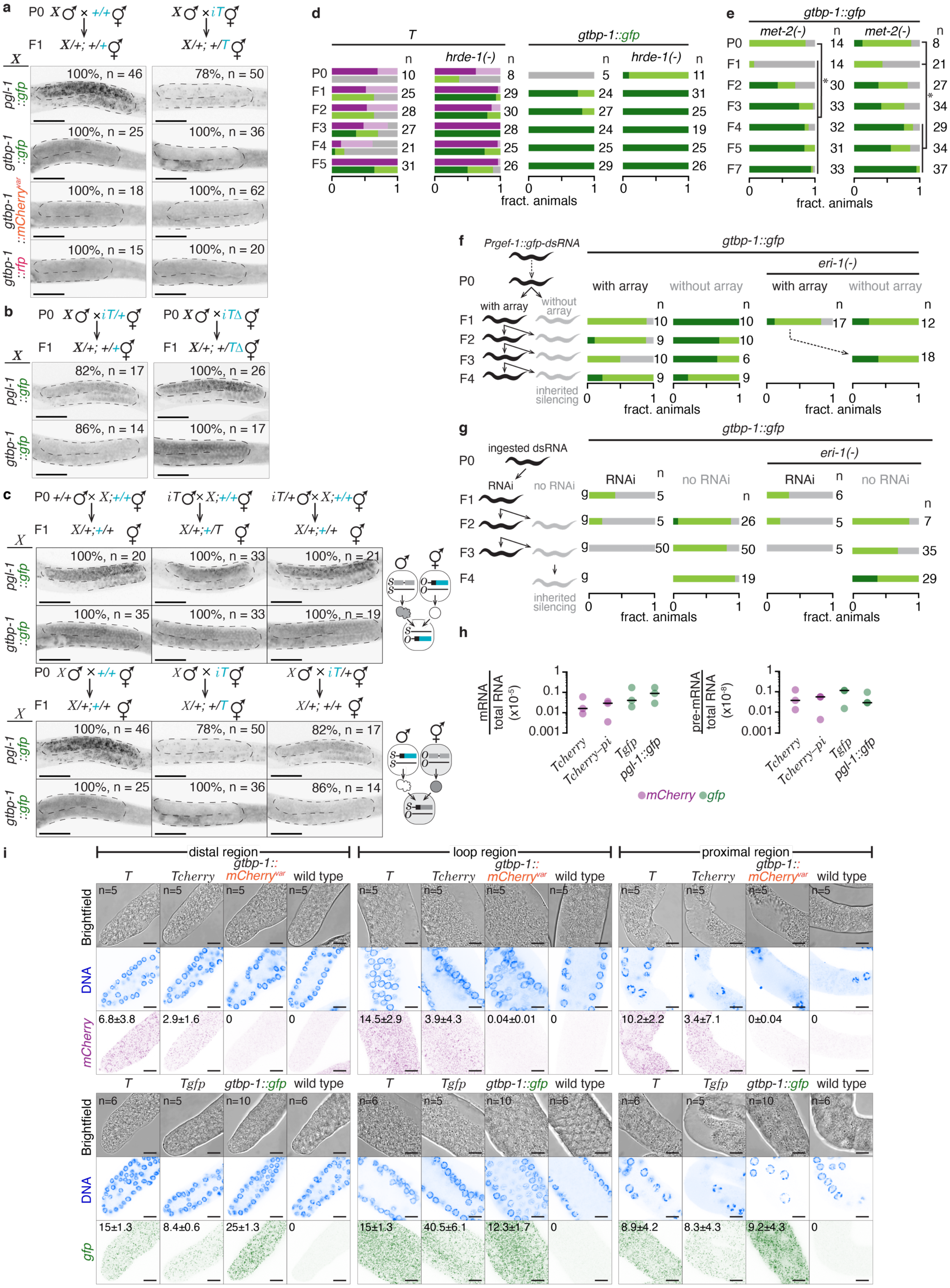
a
P0 X♂ × +/+⚥
F1 X/+; +/+⚥
X♂ × iT⚥
X/+; +/T⚥
X
pgl-1::gfp
100%, n = 46
78%, n = 50
gtbp-1::gfp
100%, n = 25
100%, n = 36
gtbp-1::mCherry^var
100%, n = 18
100%, n = 62
gtbp-1::rfp
100%, n = 15
100%, n = 20
b
P0 X♂ × iT/+⚥
F1 X/+; +/+⚥
P0 X♂ × iTΔ⚥
F1 X/+; +/TΔ⚥
82%, n = 17
100%, n = 26
86%, n = 14
100%, n = 17
c
P0 +/+♂ × X;+/+⚥
F1 X/+;+/+
iT♂ × X;+/+⚥
X/+;+/T
iT/+♂ × X;+/+⚥
X/+;+/+
100%, n = 20
100%, n = 33
100%, n = 21
100%, n = 35
100%, n = 33
100%, n = 19
P0 X♂ × +/+⚥
X/+;+/+
X♂ × iT⚥
X/+; +/T
X♂ × iT/+⚥
X/+; +/+
100%, n = 46
78%, n = 50
82%, n = 17
100%, n = 25
100%, n = 36
86%, n = 14
d
T
hrde-1(-)
gtbp-1::gfp
hrde-1(-)
n
P0 10 8 5 11
F1 25 29 24 31
F2 28 30 27 25
F3 27 28 24 19
F4 21 25 25 25
F5 31 26 29 26
fract. animals
e
gtbp-1::gfp
met-2(-)
met-2(-)
n
P0 14 8
F1 14 21
F2 30 27
F3 33 34
F4 32 29
F5 31 34
F7 33 37
fract. animals
f
Prgef-1::gfp-dsRNA
P0
with array
without array
F1
F2
F3
F4
inherited silencing
gtbp-1::gfp
eri-1(-)
with array
without array
with array
without array
n
10 10 17 12
9 10
10 6 18
9 9
fract. animals
g
ingested dsRNA
P0
RNAi
no RNAi
F1
F2
F3
F4
inherited silencing
gtbp-1::gfp
eri-1(-)
RNAi
no RNAi
RNAi
no RNAi
n
9 5 6
9 5 26 5 7
9 50 50 5 35
9 19 29
fract. animals
h
mRNA / total RNA (x10^-5)
pre-mRNA / total RNA (x10^-8)
1
0.1
0.01
0.001
Tcherry
Tcherry-pi
Tgfp
pgl-1::gfp
mCherry
gfp
i
distal region
loop region
proximal region
T
Tcherry
gtbp-1::mCherry^var
wild type
Brightfield
DNA
mCherry
n=5
6.8±3.8
2.9±1.6
0
0
14.5±2.9
3.9±4.3
0.04±0.01
0
10.2±2.2
3.4±7.1
0±0.04
0
T
Tgfp
gtbp-1::gfp
wild type
n=6
n=5
n=10
n=6
gfp
15±1.3
8.4±0.6
25±1.3
0
15±1.3
40.5±6.1
12.3±1.7
0
8.9±4.2
8.3±4.3
9.2±4.3
0

**Extended Data Figure 10. Recovery of gene expression can occur after enhanced silencing and does not correlate with transcript abundance or localization in the germline.**

**a**, Males that express homologous (*gfp*) or non-homologous (synonymous *mCherry* variant or *rfp*) sequences fused to endogenous genes ($X$ = *pgl-1* or *gtbp-1*) expressed in the germline (*pgl-1*) or ubiquitously (*gtbp-1*) were mated with non-transgenic or $iT$ hermaphrodites and fluorescence of PGL-1::GFP, GTBP-1::GFP, GTBP-1::mCherry or GTBP-1::RFP was imaged in cross progeny. **b**, Males that express *pgl-1::gfp* or *gtbp-1::gfp* were mated with hemizygous $iT$ or homozygous $iT\Delta$ hermaphrodites and GFP fluorescence from the tagged gene was scored in cross progeny that did not inherit $iT$. **c**, Animals that express *pgl-1::gfp* or *gtbp-1::gfp* were mated with homozygous or hemizygous $iT$ animals and GFP fluorescence from the tagged gene was scored in cross progeny. Germlines of representative cross progeny at L4 stage are outlined and percentages of animals with the depicted expression are indicated (**a**-**c**). **d**, Hermaphrodites expressing *T* or *gtbp-1::gfp* in a wild-type or *hrde-1(-)* background were exposed to *gfp* RNAi for 24 hours and descendants in subsequent generations (F1-F5) were scored. Animals of the same genotype exposed to control RNAi did not show any silencing of *gfp* or *mCherry*. For *mCherry* silencing in *T*, P0 expression is significantly different from F4 and F5 generations in wild-type and P0 expression is significantly different from all generations except F2 in *hrde-1(-)* background. For GFP expression from *T*, in a wild-type background P0 expression is significantly different from F1-F3 and in an *hrde-1(-)* background P0 expression is significantly different from all generations. For *gtbp-1::gfp* silencing, P0 expression is significantly different from all generations in wild-type and *hrde-1(-)* backgrounds. **e**, Animals expressing *gtbp-1::gfp* in a *met-2(-)* background (additional replicates done alongside Fig. 4d) were fed *gfp* dsRNA for a single generation and scored for GFP fluorescence as in Extended Data Fig. 1 in descendants. **f**, *gtbp-1::gfp* animals expressing neuronal dsRNA against *gfp* (*Prgef-1::gfp-dsRNA*, black) from a mitotically unstable array can have progeny with or without the array. Animals expressing dsRNA in a wild-type or *eri-1(-)* background with or without the dsRNA array were scored. **g**, *gtbp-1::gfp* animals fed dsRNA (black) for one, two or three consecutive generations and their untreated progeny in a wild-type or *eri-1(-)* background were scored. **h**, *mCherry* and *gfp* mRNA levels were measured by qRT-PCR between animals expressing $Tcherry$ or $Tcherry\text{-}pi$ and $Tgfp$ or *pgl-1::gfp* respectively. **i**, Animals that express DNA or protein sequence variants of *mCherry* (*top*) or *gfp* (*bottom*) genes were subjected to smFISH against *mCherry* or *gfp* transcripts within dissected gonads. Numbers within images refer to number of RNAs per 100 μm$^2$ with standard error of the mean. Animals with median values of fluorescence or RNA signal in the distal region are represented along with the loop and proximal regions within the same animals. Number of animals imaged per region is indicated within the brightfield image. In all panels, scoring of silencing, number of animals assayed, and blue font are as in Fig. 2a. Scale bars are 50 μm (**a-c**) or 10 μm (**i**). Asterisks indicate $P < 0.05$ from $\chi^2$ test. 'ns', statistically not significant.

#### Extended Data Tables

**Extended Data Table 1.** Reports on heritability of RNA silencing suggest that transgenerational silencing does not occur with every target gene.

| Target | | generations of inherited silencing | Reference |
|---|---|---|---|
| *dpy-11, mex-3, unc-22,*<br>*lir-1, lin-15, unc-15,*<br>*dpy-13, sqt-3, dpy-28,*<br>*pos-1, par-1,*<br>*dpy-11* | <br><br><br><br>*eri-1(-) bkgd.* | 1 | Burton et al., 2011, Winston et al., 2002, Fire et al., 1998<br>Tabara et al., 1999, Guang et al., 2010, Burkhart et al., 2011<br>Mao et al., 2015, Xu et al., 2018, Burkhart et al., 2011<br>Spracklin et al., 2017, Wan et al., 2018 |
| *Plet-858::gfp, Psur-5::sur-5::gfp*<br>*Pmyo-3::gfp, pes-10::gfp* | | 1 | Timmons et al., 2003, Xu et al., 2018,<br>Fire et al., 1998, Guang et al., 2010 |
| *Pdpy-30::mcherry::gpd-2/3::gfp* | | 1 | Sapetschnig et al., 2015 |
| *mom-2, pos-1, sgg-1,*<br>*unc-22, dpy-11* | | 2 | Grishok et al., 2000, Ashe et al., 2015 |
| *Ppie-1::gfp::H2B* | | 3 | Wan et al., 2018 |
| *oma-1* | *met-2(-); set-25(-);*<br>*set-32(-) bkgd.* | 2-5 | Buckley et al., 2012, Burton et al., 2011,<br>Houri Ze'evi et al., 2016, Spracklin et al., 2017,<br>Perales et al., 2018, Wan et al., 2018, Lev et al., 2018 |
| *Ppie-1::gfp::H2B* | | 1-9 | Buckley et al., 2012, Ashe et al., 2012, Houri Ze'evi et al, 2016,<br>Spracklin et al., 2017, Woodhouse et al., 2018, Xu et al., 2018,<br>Weiser et al., 2017 |
| *Pcdk-1::gfp* | | > 10 | Shirayama et al., 2012 |
| *Ppie-1::gfp::H2B* | *eri-1(-) bkgd.* | > 20 | Vastenhouw et al., 2006 |
| *Ppie-1::gfp::H2B* | *heri-1(-) bkgd.* | > 23 | Perales et al., 2018 |
| *oma-1*<br>*Pmex-5::gfp* | *met-2(-) bkgd.* | > 10<br>> 30 | Lev et al., 2017, Lev et al., 2018 |
| *Pmex-5::mCherry::gfp* | | > 25 | Devanapally et al., 2015 |

**Extended Data Table 2.** Comparison of mating-induced silencing with related epigenetic phenomena.

| Phenomenon | Reference(s) for the phenomenon | Similarity with mating-induced silencing | Difference from mating-induced silencing |
|---|---|---|---|
| Paramutation in plants, flies, or mice | 61, 62, 63, 61, 65 | Silencing is transgenerational. Silenced allele inherited through either gamete can silence homologous sequences. | Silencing cannot be predictably initiated. When a silenced allele induces meiotically heritable silencing of another allele, this allele also becomes a silencing allele. |
| RNA induced epigenetic silencing (RNAe) | 18, 19, 20, 69, 72 | Initiation requires PRG-1; maintenance requires HRDE-1. Silencing is transgenerational. | Silencing cannot be predictably initiated. The same DNA inserted into the same locus can show expression or silencing. Changes upon mating, if any, are unknown. |
| Multi-generational RNAe caused by meiotic silencing by unpaired DNA | 77 | Initiation requires PRG-1. *oxSi487* (*T* in our study) introduced through the male parent showed silencing in cross progeny. | Effect of introducing *oxSi487* through the hermaphrodite parent on silencing in cross progeny or its hemizygous descendants was not tested. |
| RNA-induced epigenetic gene activation (RNAa) | 30, 36 | Extragenic signal can be inherited from male to control gene expression in progeny. Inheritance of an active transgene from hermaphrodite affects expression of paternally inherited transgene. | Extragenic signals inherited from sperm promote expression. |
| Meiotic silencing by unpaired DNA | 78 | Silencing of DNA is epigenetic. | DNA must be upaired during meiosis for silencing. |
| Epigenetic licensing of *fem-1* | 27 | Maternal transcript of a gene is sufficient to enable expression of the paternal copy in the zygote. | Repeated crossing was required for increased severity of silencing. |
| Genomic imprinting and parent of origin effects | 38, 70, 80 | Silencing occurs when a gene is inherited through a specific gamete. | Expression is reset upon passage through the other gamete. |
| Transposon silencing in flies | 65, 81 | Inherited piRNAs silence a paternally inherited gene. | Maternal transcript does not prevent gene silencing. |
| Transvection in flies | 82 | Interaction between alleles on homologous chromosomes can result in changed expression. | Changes in gene expression are not heritable. |
| Licensing by DNA sequences | 45 | Not all transgenes are susceptible to germline silencing. | Initiation of silencing is independent of mating. |

**Extended Data Table 3.** Reagents used for Cas9-mediated genome editing.

| Allele name | CRISPR edit | Primers used to make: | | Length of homology repair template | Concentration of reagents used (pmol/µl) | | | | |
|---|---|---|---|---|---|---|---|---|---|
| | | DNA template for sgRNA transcription or crRNA sequence | Homology repair dsDNA or ssDNA template | | First sgRNA/ crRNA | Second sgRNA/ crRNA | Homology repair template | *dpy-10* sgRNA#/ crRNA | *dpy-10* homology repair template |
| *+* | *dpy-10(-)* in wild type | P57 (FOR), P43 (REV) | P58 (ssDNA) | 100 b | - | - | - | 3.05 | 0.66 |
| *T* | *dpy-10(-)* in *oxSi487* | P57 (FOR), P43 (REV) | P58 (ssDNA) | 100 b | - | - | - | 3.05 | 0.66 |
| *T** | *mCherry* mutation in *oxSi487* $ | P64 (FOR), P43 (REV), P163 (FOR) | Left: P65 + P66, Right: P67 + P68, Fusion: P69 + P70 | 309 bp | 1.6 | 1.4 | 0.12 | 1.3 | 0.66 |
| *TΔ** | *mCherry* mutation in *jamSi19* (*TΔ*) | P46 (FOR), P43 (REV) | P50 (ssDNA) | 60 b | 6.05 | - | 8.85 | 3.05 | - |
| *TΔΔ** | *mCherry* mutation in *jamSi25* (*TΔΔ*) | P46 (FOR), P43 (REV) | P50 (ssDNA) | 60 b | 6.05 | - | 8.85 | 3.05 | - |
| *TΔ* | Deletion of *gfp* and *tbb-2 3' utr* from *oxSi487* | P59 (FOR), P43 (REV) | Left: P60 + P61, Right: P62 + P52, Fusion: P63 + P54 | 1074 bp | 2.96 | - | 0.08 | 3.05 | 0.66 |
| *TΔΔ* | Deletion of *Punc-119* from *jamSi19* (*TΔ*) | P55 (FOR), P43 (REV) | P56 (ssDNA) | 60 b | 8.4 | - | 1.53 | 8.16 | 1.52 |
| *TΔΔΔ* | Deletion of *h2b* from *jamSi25* (*TΔΔ*) | P42 (FOR), P43 (REV) | Left: P44 + P45, Right: P47 + P48, Fusion: P80 + P81 | 1604 bp | 11.16 | 12.87 | 0.31 | 2.89 | 0.62 |
| *Tcherry-pi N* | Deletion of *mCherry C-terminus* (*Tcherry-pi*) | P164 (crRNA), P165 (crRNA) | P168 (ssDNA) | 70 b | 4.0 | 4.0 | 24 | 2.4 | 100 |
| *Tcherry-pi C* | Deletion of *mCherry N-terminus* (*Tcherry-pi*) | P166 (crRNA), P167 (crRNA) | P169 (ssDNA) | 70 b | 18.6 | 11.2 | 24 | 2.4 | 100 |
| *Tcherry-pi exon4* | Deletion of three *mCherry* exons from *jamSi37* (*Tcherry*) | P173 (crRNA), P174 (crRNA) | P175 (ssDNA) | 70 b | 4.9 | 4.9 | 3.6 | 2.4 | 100 |
| *T-orf* | Deletion of *mCherry* ORF from *jamSi37* (*Tcherry*) | P170 (crRNA), P171 (crRNA) | P172 (ssDNA) | 70 b | 4.8 | 4.8 | 25 | 2.4 | 100 |
| *iT* | *dpy-2(-)* repair in *iT dpy-2(-)* | P42 (FOR), P43 (REV) | P101 (ssDNA) | 60 b | 7.2 | - | 0.6 | - | - |
| *rde-8(-)* | *rde-8* mutation in *iT* | P100 (FOR), P43 (REV), P102 (FOR) | P103 (ssDNA) | 60 b | 8.1 | 10.9 | 13.5 | 6.9 | 6.5 |
| *set-32(-)* | *set-32* mutation in *iT* | P104 (FOR), P43 (REV), P105 (FOR) | P106 (ssDNA) | 60 b | 3.9 | 3.9 | 7.5 | 2.8 | 7.5 |
| *heri-1(-)* | *heri-1* mutation in *iT* | P107 (FOR), P43 (REV), P108 (FOR) | P109 (ssDNA) | 60 b | 3.7 | 3.7 | 7.5 | 2.3 crRNA (P110) 2.7 tracrRNA (P111) | 7.5 |

$ refers to cases where the resulting edit was not the originally intended edit and therefore does not relate to the reagents injected.
### *dpy-10* sgRNA was *in-vitro* transcribed using a DNA template generated using primers P57 (forward) and P43 (reverse).
